# Functional regulatory variants implicate distinct transcriptional networks in dementia

**DOI:** 10.1101/2021.06.14.448395

**Authors:** Yonatan A. Cooper, Jessica E. Davis, Sriram Kosuri, Giovanni Coppola, Daniel H. Geschwind

## Abstract

Predicting functionality of noncoding variation is one of the major challenges in modern genetics. We employed massively parallel reporter assays to screen 5,706 variants from genome-wide association studies for both Alzheimer’s disease (AD) and Progressive Supranuclear Palsy (PSP). We identified 320 functional regulatory polymorphisms (SigVars) comprising 27 of 34 unique tested loci, including multiple independent signals across the complex 17q21.31 region. We identify novel risk genes including *PLEKHM1* in PSP and *APOC1* in AD, and perform gene-editing to validate four distinct causal loci, confirming complement 4 (*C4A*) as a novel genetic risk factor for AD. Moreover, functional variants preferentially disrupt transcription factor binding sites that converge on enhancers with differential cell-type specific activity in PSP and AD, implicating a neuronal *SP1*-driven regulatory network in PSP pathogenesis. These analyses support a novel mechanism underlying noncoding genetic risk, whereby common genetic variants drive disease risk via their aggregate activity on specific transcriptional programs.

**One Sentence Summary:** High-throughput functional analysis of GWAS loci reveals cell-type specific regulatory networks that mediate genetic risk for dementia.

## Main Text

Neurodegenerative disorders such as Alzheimer’s disease (AD) and Progressive Supranuclear Palsy (PSP) are a major and growing cause of morbidity and mortality worldwide, with AD alone expected to impact 135 million individuals by 2050 (*1*). Given the lack of any disease modifying therapeutics, there is a significant need for investigation of causal disease mechanisms. Sporadic AD and PSP, both known as tauopathies because of the pathological deposition of tau in the brains of affected individuals (*2*), are complex polygenic traits with heritability estimates of between 40-80% (*3, 4*). Over the last decade a number of genome-wide association studies (GWAS) have identified numerous susceptibility loci (*5–12*). However, most of these loci fall in noncoding regions of the genome and encompass numerous variants due to linkage disequilibrium (LD), which has hampered the identification of underlying regulatory variants and associated risk genes (*13–15*). This has posed a particular challenge for the interpretation of extended complex haplotypes harboring multiple independent association signals such as the H1 pan-neurodegenerative risk haplotype at 17q21.31 that includes the *MAPT* locus (*12, 16, 17*), and the AD risk locus at 19q13.32 that harbors *APOE* (*18–20*).

Although a number of statistical fine-mapping approaches have been developed to identify causal GWAS variants, these methods perform poorly on underpowered datasets or regions of extended LD (reviewed in (*21*)). Similarly, prioritization algorithms that score variant pathogenicity by leveraging features such as evolutionary conservation and chromatin annotations underperform in noncoding regions of the genome, or are nonspecific (*22, 23*). It is becoming increasingly recognized that functional methods are necessary to identify true causal variants within most loci, but the sheer numbers of variants challenge most experimental approaches. Massively parallel reporter assays (MPRA) provide a solution, enabling high-throughput experimental characterization of the transcriptional-regulatory potential of noncoding DNA elements (*24–27*). In an MPRA, many regulatory elements are cloned into an expression vector harboring a reporter gene and a unique DNA barcode to create an expression library that is assayed via high-throughput sequencing (*24*). MPRAs have prioritized functional common variation for lymphoblastoid eQTLs (*28*), red blood cell traits (*29*), cancer (*30*), adiposity (*31*), and osteoarthritis (*32*). However, they have not been systematically applied to neurodegeneration or any neurologic disorder.

In this study, we apply MPRA to characterize common genetic variation associated with two distinct neurodegenerative disorders that both share tau pathology, AD and PSP. We screen 5,706 unique variants comprising 34 genome-wide significant or suggestive loci derived from three GWASs (*6, 7, 10*), including a comprehensive assessment of disease-associated variants within the 17q21.31 and 19q13.32 regions. We identify 320 variants with significant allelic skew at a conservative false discovery rate and implicate novel risk genes including *PLEKHM1* in PSP and *APOC1* and *C4A* in AD. We find that SigVars preferentially disrupt transcription factor binding sites to impact specific transcriptional programs, and highlight a neuronal SP1 transcriptional network underlying PSP risk. Furthermore, using genome editing we validate rs13025717, rs636317, rs6064392, and rs9271171 as functional variants regulating *BIN1, MS4A6A, CASS4*, and *HLA-DQA1/C4A* expression, respectively. Finally, we provide evidence that MPRA-determined variant prioritization is discordant with existing computational algorithms, underscoring the utility of high-throughput experimental datasets.

## Results

### MPRA to identify candidate regulatory GWAS variants

We conducted a staged analysis to identify regulatory variants underlying GWAS loci for two neurodegenerative tauopathies – Alzheimer’s disease (AD) and Progressive Supranuclear Palsy (PSP) – using massively parallel reporter assays (MPRA) (Fig. 1a). In stage 1 (MPRA 1), we identified all variants in linkage disequilibrium (LD; r^2^ > 0.8) with the 1,090 genome-wide significant (p < 5×10^-8^) variants from an AD GWAS (*6*) and 3,626 genome-wide significant variants from a PSP GWAS (*10*). After filtering for bi-allelic noncoding variants, this resulted in a list of 5,223 variants encompassing 14 AD and 5 PSP GWAS loci. Both alleles of each variant were centered in 162 base pairs (bp) of genomic context, synthesized as 210 bp oligonucleotides, and cloned into a custom MPRA vector along with degenerate barcodes to create an expression library (Supplemental Fig. 1; Methods). In stage 2 (MPRA 2), we sought to replicate 326 variants screened in MPRA 1 to asses reproducibility, test the importance of oligo configuration on assay performance, and screen an additional 483 variants encompassing 11 new loci from 2 recent AD GWAS (*6, 7*) and 4 new suggestive loci for PSP (*10*) (Fig. 1a; Table 1; Methods).

**Figure 1:**
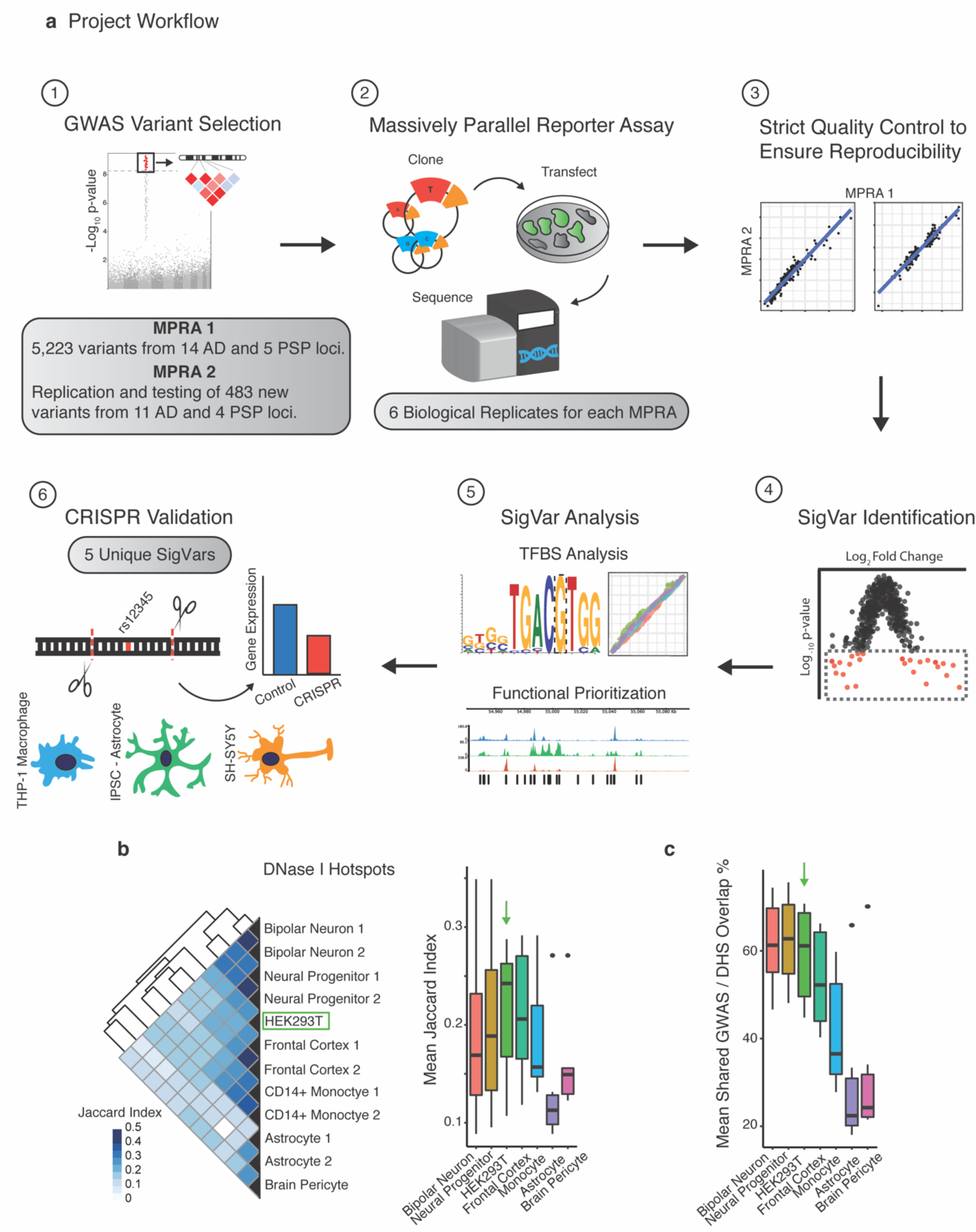
(**A**) Project workflow: 1) 5,223 genome-wide significant variants and LD partners encompassing 14 AD and 5 PSP GWAS loci were selected in MPRA 1. For MPRA 2, select variants identified in MPRA 1 were replicated. An additional 483 variants from 11 AD and 4 PSP loci were also tested. 2) Both alleles of each variant were barcoded and cloned into an expression library that was transfected into HEK293T cells. Allele expression was quantified by next-generation sequencing of associated barcodes. 3) Strict quality control was performed to confirm within-assay and between-experiment reproducibility. 4) Comparison of expression between alleles enabled identification of variants with significant allele-specific transcriptional skew (SigVars). 5) SigVars were used for transcription factor binding site-disruption analysis and benchmarking of computational prediction algorithms. SigVars were further prioritized using brain-specific genomic annotations and 6) five top variants were selected for validation using CRISPR/Cas9 genome-editing in brain-relevant cell lines (THP-1, IPSC-astrocytes, SH-SY5Y). (**B-C)** comparison of the chromatin landscape between divergent brain cell types. (**B**) left – Heatmap quantifying overlap (Jaccard index) of DNase hotspots (ENCODE project) between primary brain cell types and HEK293T cells. Right – boxplot displaying the mean pairwise Jaccard index for each cell type. (**C**), GWAS variants tested in this study were overlapped with DNase hotspots from each cell type. The boxplot displays the mean pairwise probability that a GWAS variant within open chromatin in a given cell type also overlaps open chromatin in another cell type. Green arrows highlight HEK293T. Error bars = S.E.M.

**Table 1.**
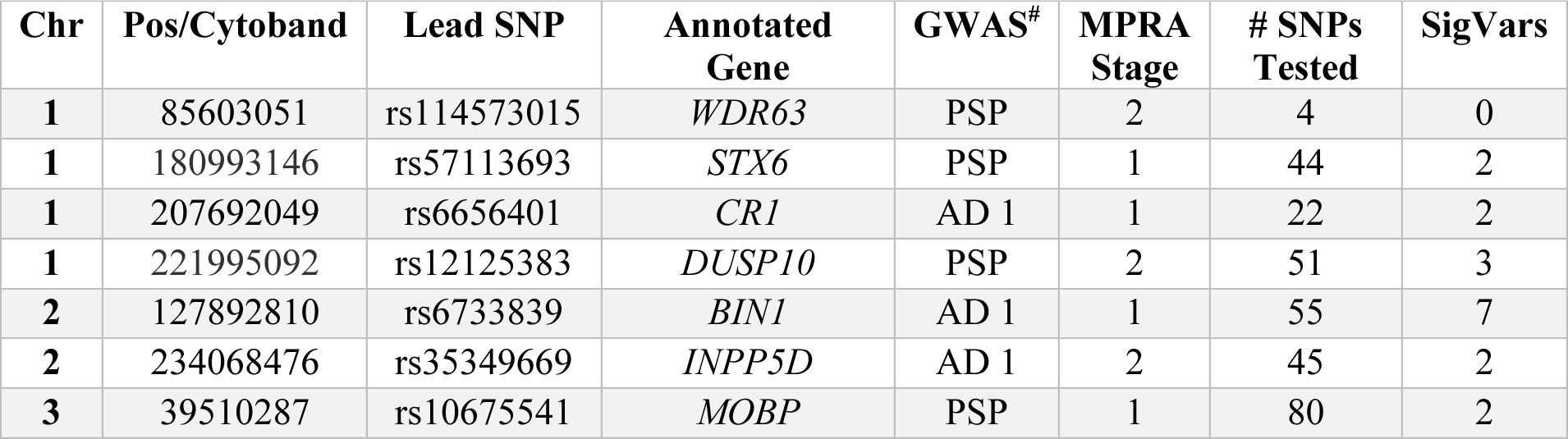
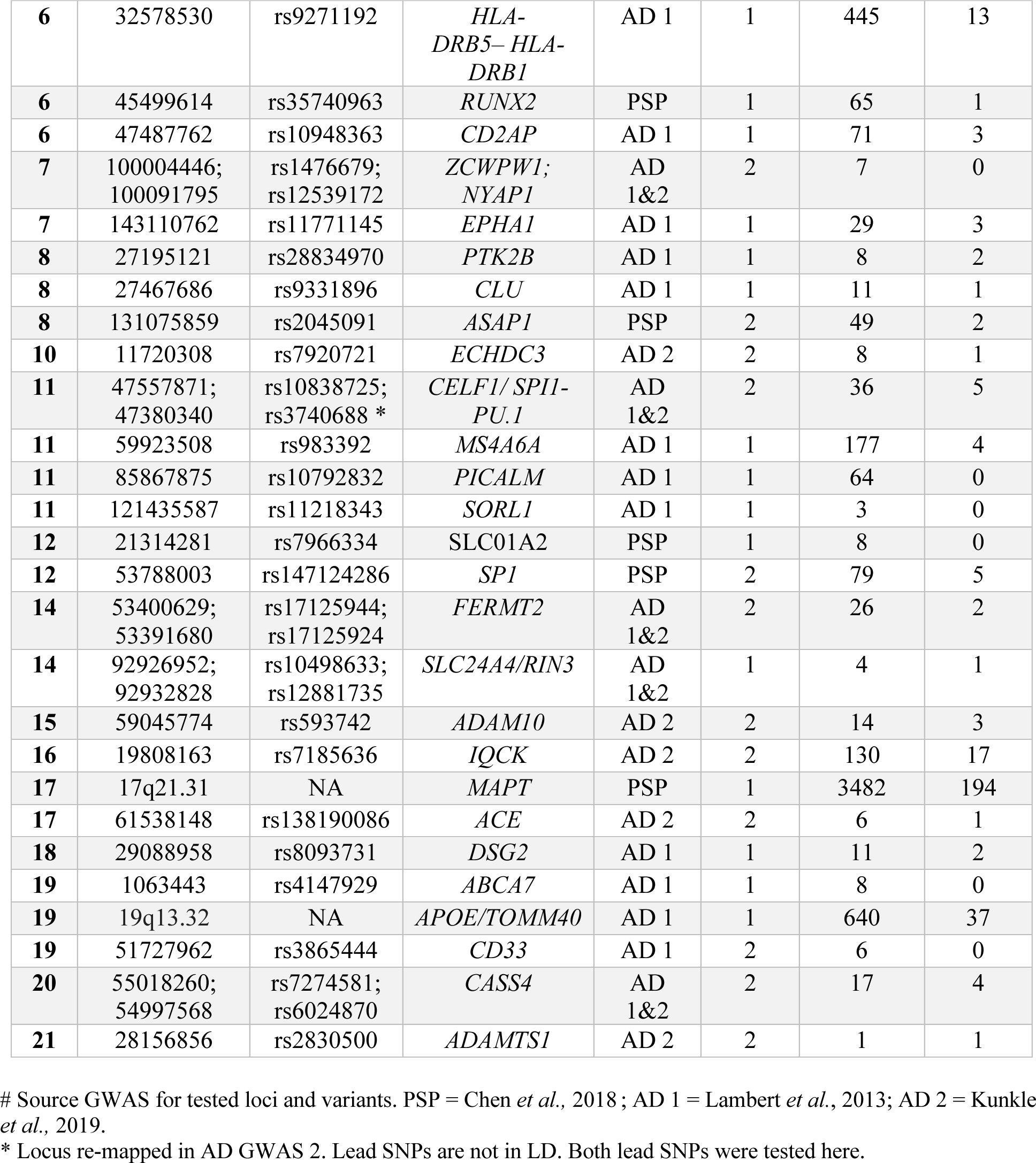
AD and PSP GWAS Loci. Description of GWAS loci and variation tested in this study. Shown are the locus lead SNPs and annotated genes as described in the listed GWASs. For each locus: the MPRA stage in which it was tested, the number of variants tested per locus, and the number of variants with significant allelic skew (FDR q < 0.01; SigVars) ultimately identified.

The performance of MPRAs to detect allelic skew depends upon high library transfection efficiency (*33*), necessitating the use of easy to transfect cell lines, which in published studies have included HEK293T, K562, and HepG2 cells, among others (*28, 34*). However, AD and PSP disease risk variants fall within open chromatin across several different neuronal and glial cell types, many of which are non-overlapping (*7*). Using available ENCODE DHS data (*35*), we found poor DNA accessibility overlap between divergent brain cell types, such as astrocytes and neural progenitors (mean Jaccard index = 0.14). Notably, HEK293T cells had the highest mean pairwise Jaccard index (0.22; Fig. 1b) compared with all brain cell types and tissues, which was not driven by a larger number, or increased width of peaks in HEK293T cells. Moreover, we found that AD and PSP GWAS variants that fell within open chromatin in any brain cell type were also likely to fall within open chromatin in HEK293T cells (mean = 60%; Fig. 1c). These data indicated that HEK293T cells would provide an optimal model for such high-throughput screening in a single cell line, and they were chosen for the MPRA.

We performed MPRA (n = 6 biological replicates), obtaining activity measurements from at least 5 unique barcodes for both alleles for 4,732 of 5,223 variants (∼91%), with a median library complexity of 40 barcodes/allele (Fig. S2a). Intra-library reproducibility was high; allele correlation between replicates had mean Pearson’s r = 0.95 (Fig. 2b) and normalized barcode correlations were consistent with expectations (r = 0.55; Fig. S2c). Overall, we observed that ∼19% of library elements were transcriptionally active in our assay (Fig. 2c), in concordance with previous estimates (*28, 29*).

**Figure 2:**
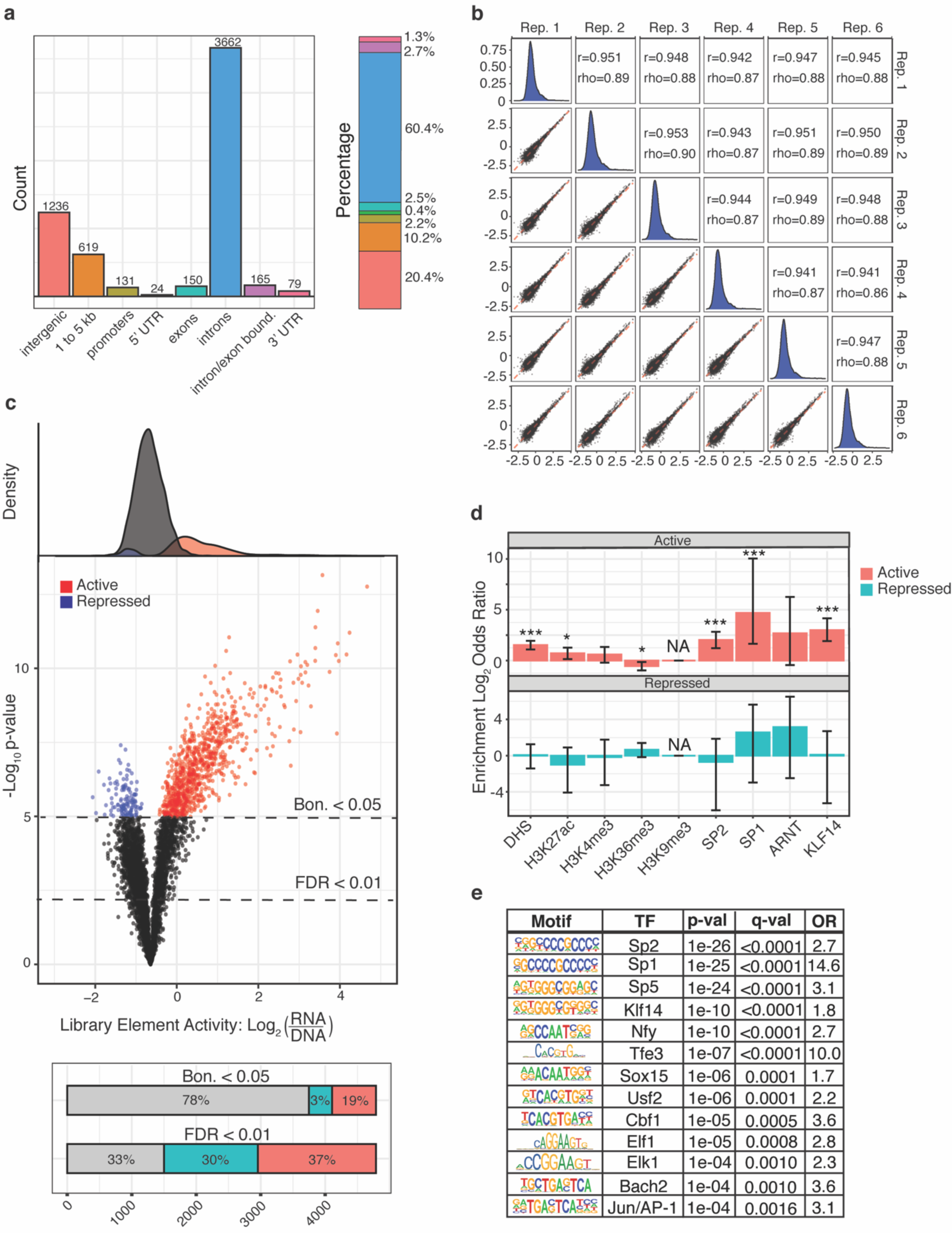
The MPRA exhibited strong technical performance with active MPRA elements enriched for functional chromatin annotations. (**A**) The bar charts (counts and percentages) show genomic annotations for 5,223 variants tested in MPRA 1. (**B**) MPRA 1 exhibits high reproducibility between biological replicates (mean r = 0.95; n = 6). Panels show inter-replicate, pairwise correlations of median log_2_ normalized barcode counts for all alleles passing filter (n = 9,464). Red line = regression line of best fit, Pearson’s correlation, all p < 2×10^-16^. (**C**) Identification of active and repressed MPRA elements. MPRA library transcriptional activity was quantified by comparing the median normalized barcode count for each element against the median of the whole library (n = 6 replicates; one-sample Mann-Whitney-U test; two-sided, Bonferroni correction). Significantly (Bonferroni p < 0.05) increased (active) and decreased (repressed) library elements highlighted on the volcano plot in red and blue respectively. (**D-E**) Active elements are enriched for relevant chromatin features and TFBSs. (**D**) Enrichment log_2_ odds ratios (Fisher’s exact test) of active and repressed elements within HEK293T ChIP-seq peaks for both histone and TF marks. Error bars = 95% CI, *** FDR-adjusted q < 0.001 (BH method), * q < 0.05. (**E**) Active elements enrich for predicted TF binding motifs, including *SP*/*KLF*, and *FOS/JUN* family TFs.

We next assessed the functional genome annotations associated with active versus non-active elements. Using available ENCODE data for HEK293T cells, we found depletion of the H3K36me3 mark in active elements (Fisher’s exact test; log_2_ OR = -0.54, FDR-adjusted q = 0.01). Conversely, active elements were highly enriched for DHS sites (log_2_ OR = 1.52, q = 1×10^-10^), which are indicative of accessible chromatin. Furthermore, H3K27ac and H3K4me3 marks, delineating active enhancers and promoters respectively, were likewise enriched in active elements (Fig. 2d; Fig. S2d). ChIP-seq peaks for specific Transcription Factors (TF) were similarly enriched in active elements (Fig. 2d; Fig. S2d). We also assessed for enrichment of specific TFBSs within active elements and identified significant enrichments of *SP*/*KLF*, *ETS*, and *AP-1* family members (Fig. 2e).

We applied a stringent statistical threshold to identify variants with significantly different transcriptional efficacy between alleles, termed *SigVars* (two-sided Mann-Whitney-U test, FDR q < 0.01, Benjamini-Hochberg method), identifying 267 SigVars. We next analyzed the second MPRA (n = 6 biological replicates), which maintained high quality with median library barcode complexity of 54 and high allele level (mean r = 0.99) and barcode level (mean r = 0.84) correlations between replicates (Fig. S3a-b). Assessments of reproducibility between the separate MPRA experiments also revealed that our assay was robust; the correlations between activity scores (Pearson’s r = 0.98, p < 2×10^-16^) as well as effect sizes (r = 0.94, p < 2×10^-16^; Fig. 3a) for replicated variants were both high, and 152 of 186 (82%) re-tested SigVars were reproduced in the second MPRA (replication Bonferroni p < 0.05). Placing oligos in the reverse orientation completely abolished reporter activity as expected (Fig. S3c-d; Supplementary Text).

**Figure 3:**
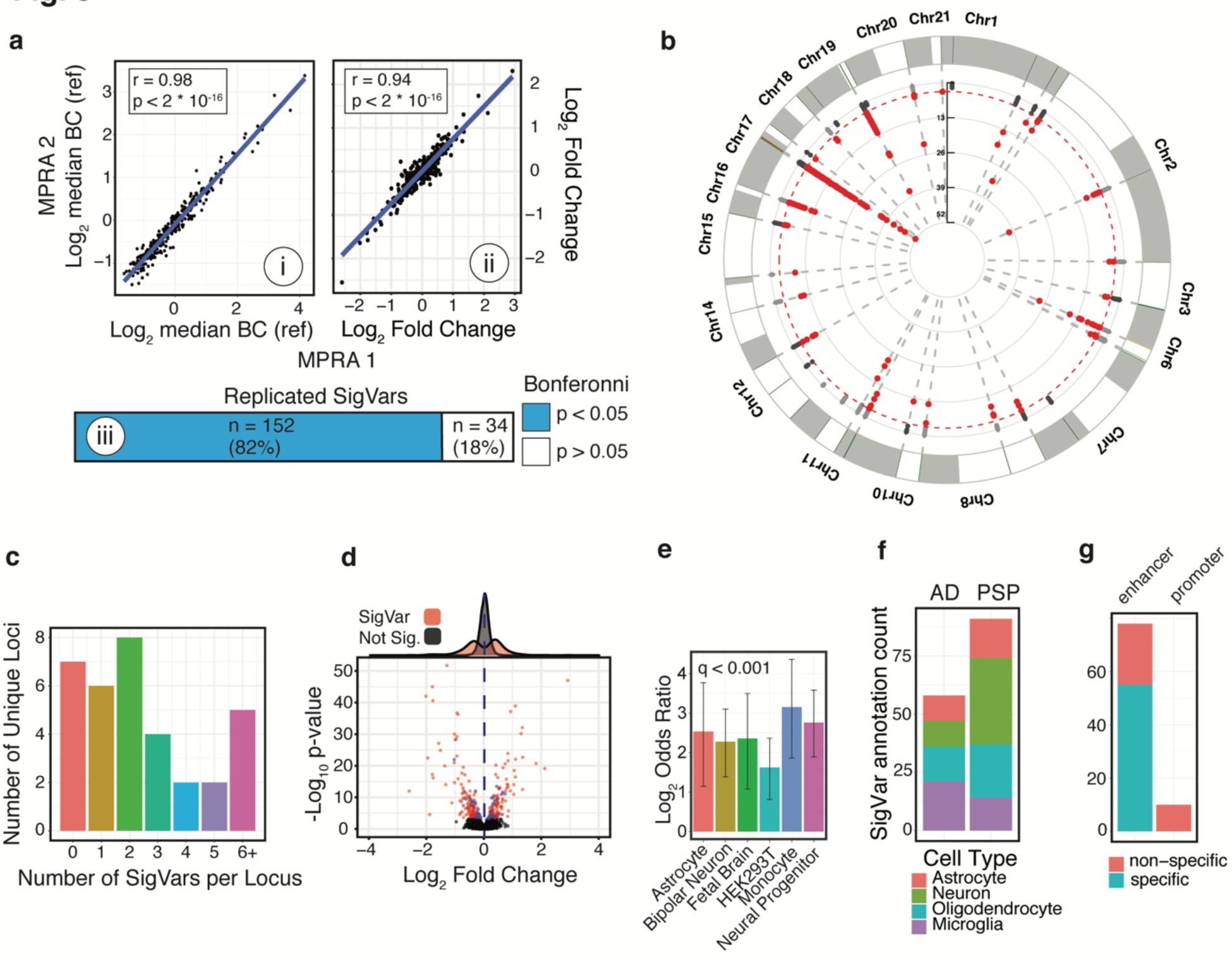
Identification of variants with significant allelic skew (SigVars) from both MPRA stages. (**A**) Reproducibility of MPRA across experimental stages. 326 variants, including 186 SigVars identified from the preliminary MPRA (1) were re-tested in a follow-up MPRA (2). i) Reference allele transcriptional activity and ii) log_2_ effect sizes (alt/ref allele) show strong correlation (Pearson’s r = 0.98, 0.94, p < 2×10^-16^) between experiments. BC = barcode count. iii) 152 of 186 assessed SigVars from MPRA 1 were replicated in MPRA 2 at replication Bonferroni p < 0.05. (**B**) Manhattan plot of 5,340 unique variants successfully tested across both experiments. Red indicates SigVars at FDR-adjusted q < 0.01 (BH method). (**C**) Histogram of the number of SigVars identified per GWAS locus LD block (median = 2). (**D**) Volcano plot shows log_2_ allelic skew effect sizes and -log_10_ p-values for 5,340 unique variants tested by MPRA. SigVars for MPRA stage 1 (red) and 2 (blue) highlighted. blue line = median effect size. (**E**) Enrichment log_2_ odds ratios (Fisher’s exact test) of SigVars within DHSs of various brain cell types (ENCODE project). Error bars = 95% CI, All FDR-adjusted q < 0.001. (**F-G**) SigVars were separated into those derived from AD or PSP GWAS and annotated for overlap with enhancer marks from sorted brain tissue (annotations (*36*)). Bar plot shows cell-type enhancer annotation counts for overlapping SigVars separated by disease. A plurality of AD SigVars fell within microglial enhancers, while PSP SigVars fell within neuronal enhancers. (**G**) most SigVars overlap enhancers present in only one cell type.

Given the high correspondence between both MPRA experiments, we combined them, obtaining activity measurements from 5,340 of 5,706 (93.6%; Supplemental Data) assayed variants and identifying 320 unique SigVars distributed across 17 chromosomes (6.0%; Fig. 3b). We identified SigVars in 27 of 34 (79%) tested GWAS loci with a median of 2 SigVars per locus (Fig. 3c; Table 1). As expected, effect sizes (alt/ref allele) were generally modest (Fig. 3d), with mean absolute SigVar log_2_ fold change of 0.53, consistent with prior work (*29*). SigVars were highly enriched within library elements that were transcriptionally active (Fig. 2c) in our screen (OR = 6.2; p < 2×10^-16^; Methods) and were also significantly enriched within DHSs within major brain cell types including astrocytes and neural progenitors, as well as monocytes, which are from the same lineage as microglia in the brain (Fig. 3e). However, when we separated SigVars derived from AD vs. PSP GWAS loci and identified those that were found to overlap enhancer marks from human brain neurons, microglia, astrocytes, and oligodendrocytes, we saw that the cell types impacted by each disorder were distinct (Methods) (*36*). A plurality of AD SigVars fell within microglial enhancers. In contrast, a plurality of PSP SigVars fell within neuronal enhancers, in concordance with recent estimates of cell type specific enrichment in SNP-based heritability for these two disorders (*37*) (Fig. 3f). Interestingly, combined across both disorders, 55/78 (71%) of these variants overlapped with cell-type specific enhancer annotations (Fig. 3g).

### Comparing computational predictions with empirically determined function

The use of computational methods for *a priori* identification of functional classes of variants is an ongoing area of active research and several predictive methods have been developed (*38*). We sought to determine the extent to which these methods recapitulate our empirical MPRA data and subsequent variant annotations, as well as previously published MPRA data in a different cell type from Tewhey *et al.* (*28*). We compared MPRA-determined effect sizes and allelic skew significance levels - derived from this study and the previously published MPRA dataset (*28*) - with regulatory scores from four widely used algorithms: CADD, CATO, GWAVA, and LINSIGHT (*39–42*). For both MPRA datasets, representing vastly different disorders and tissue types, we observed that these computational methods indeed were able to capture enrichment of regulatory signal when comparing the top vs bottom 5^th^ percentile of variants as ranked by allelic skew p-values (Fig. S4a-b). However, if we applied a statistical threshold that would identify variants with functional effects (FDR-corrected thresholds of 0.01; Methods), overall regulatory predictions were highly discordant and not strongly predictive (max AUC = 0.55, 0.56; Fig. 4a-b). MPRA effect sizes also failed to correlate with algorithm predictions, with the exception of CATO (Fig. S4c-d). We also calculated pairwise overlap metrics and found that MPRA SigVar experimental outcomes had little overlap with computational predictions (mean Cohen’s Kappa = 0; Fig. 4c-d; Methods). Interestingly, prediction algorithms had little concordance among themselves; overall mean Kappa between all predicative algorithms was 0.11 for variants in our dataset and 0.14 for variants from the other published MPRA (*28*).

**Figure 4:**
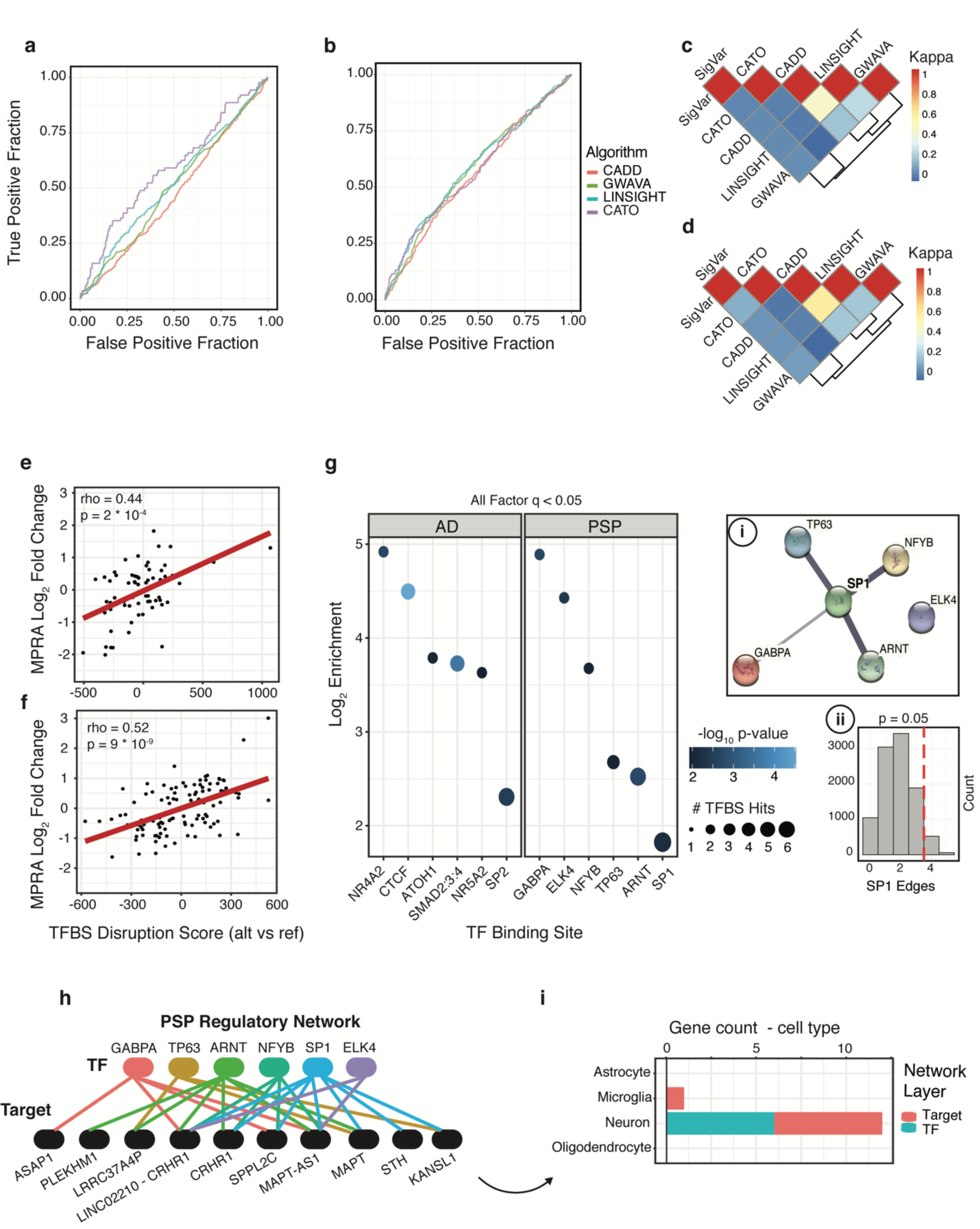
MPRA variant effect sizes correlate with TFBS disruption, but are poorly predicted by computational algorithms. (**A**) ROC curves highlight the poor predictive performance (max AUC = 0.55) of four algorithms used to score variant functionality (LINSIGHT, CADD, GWAVA, and CATO) benchmarked against MPRA SigVars (defined at q < 0.01, BH method). (**B**) same as (**A**) except using SigVars from a published MPRA dataset (*28*) (max AUC = 0.56). (**C-D**) Heatmaps show pairwise Cohen’s Kappa for MPRA SigVars and top-ranked variants scored by each algorithm (Methods) for this study (**C**) and the published study (*28*) (**D**). (**E-F**) MPRA variant effect sizes correlate with predicted TFBS-disruption scores for our dataset (**E)** and the replication dataset (**F**). Red line = OLS regression, Spearman’s rho. (**G**) Shows the log_2_ enrichments for significantly disrupted TFBSs (FDR q < 0.05, BH method), with SigVars from AD and PSP analyzed separately (color = -log_10_ enrichment p-values, size = # disrupted TFBSs; SNPS2TFBS (*43*)). Inset: i) protein-protein interaction network from the STRING (*77*) database for significantly disrupted PSP-TFs. Line thickness = strength of evidence. ii) Empirical distributions for expected PPI networkSP1 - connectivity generated by resampling (Methods). Red lines = observed PSP-TF network edges. (**H**) Network of TFs disrupted in PSP and their target genes (genes within 10 kb of disrupted TFBSs). (**I**) Bar plot counts top-expressing brain cell-type for each gene in (**H**) colored by network layer (cyan = TF, red = target gene).

### MPRA SigVars enrich for transcription factor binding site disruption

Because active MPRA elements enrich for TFBSs, we hypothesized that functional variants defined by our SigVars would be enriched for variants that disrupt TF-binding as a class. Therefore, we ran our variants through the SNPS2TFBS algorithm (*43*), which predicts TFBS disruption, finding that SigVars were enriched for variants that disrupt TFBSs (OR = 1.4, p = 0.003). We observed a similar magnitude of enrichment of TFBS-disrupting variants amongst SigVars from the published dataset from Tewhey *et al.* (*28*) (OR = 1.9, p = 8.7 ×10^-8^), and these findings were further reproducible using an alternative TFBS-scoring method (motifbreakR; Methods). Furthermore, the magnitude and directionality of predicted TFBS disruption correlated with MPRA effect sizes in both our dataset (Spearman’s rho = 0.44, p = 2.3 ×10^-4^) and the published dataset (rho = 0.52, p < 8.7 ×10^-9^; Fig. 4e-f).

Given this enrichment, we asked whether TFBS disruption alone predicts MPRA allelic skew. To do so, we scored all variants from the previously published dataset (*28*) (assay performed in K562 cells) for TFBS disruption (Methods) and found that only a proportion of TFBS-disrupting variants were also MPRA SigVars (Positive Predictive Value = 0.14). Predictive performance somewhat improved after filtering for strong disruption of TFBSs for the top 200 K562-expressed TFs (PPD = 0.22; Methods), on par with the top-performing algorithm, CATO. These findings demonstrate the importance of accounting for cell-type specific *trans-*factors for variant functional predictions, although predicted TFBS disruption only explains a subset of MPRA-defined functional variants. Interestingly, TFBS disruption was more predictive for some TFs over others. For example, disruption of *ELK1*, *ELF2*, and *GABPA* TFBSs (all ETS-family TFs), as well as SP-family TFBSs, were highly predictive of allelic skew captured by MPRA (Fig. S5), while disruption of other TF families was not.

### Enrichment of functional risk variants within disease-specific transcriptional networks

We next assessed whether SigVars were enriched within binding sites for specific transcription factors. Importantly, we determined that TFBSs disrupted by risk variants differed by disease. In AD, *NR4A2* (log_2_ OR = 4.9, p = 0.002), *NR5A2* (log_2_ OR = 3.6, p = 0.01), *ATOH1* (log_2_ OR = 3.8, p = 0.009), *SP2* (log_2_ OR = 2.3, p = 0.008), and *SMAD-*family *(SMAD2,3,4* heterotrimer, log_2_ OR = 3.7, p = 0.0002) binding sites were enriched for disrupting risk variants (all FDR-adjusted q < 0.05). Interestingly, PSP showed a different pattern of TFBS enrichment, in which five of the six TFs predicted to be enriched for binding site disruption physically interact with the transcription factor SP1 (including SP1 itself; Fig. 4g). Importantly, while these TFs are highly expressed in HEK293T cells, we do not find a general relationship between relative TF abundance and MPRA allelic skew (p > 0.05; Spearman’s correlation; Methods), indicating that expression levels do not explain our results. We also assessed enrichments at a lower statistical threshold (TFBS enrichment q < 0.1), finding a number of ETS-family TFs and SP1 interactors to be enriched within functional PSP risk variants (Fig. S6).

Further analysis indicated that the PSP-enriched TFs form a significant protein-protein interaction network with SP1 (permutation p = 0.05; Fig. 4g inset; Fig. S6; Methods), consistent with SP1’s multimerization capabilities and its activity as a core component of a broad array of gene regulatory complexes that regulate tissue specific gene expression (*44*). Interestingly, 11.1% of annotated PSP SigVars are predicted to participate specifically within this network (20% when including factors from the expanded network). Of note, the coding region for *SP1* itself falls within a suggestive PSP risk locus approaching genome-wide significance (locus combined p-value = 4.1×10^-7^) (*10*). These data, including identification of five functional regulatory variants at the *SP1* locus, provide strong evidence for disruption of an SP1-based signaling network in PSP pathophysiology.

To explore this further, we next generated a two-layer directed network composed of these six PSP TFs and their likely targets, defined as genes within the *cis*-regulatory window of TFBS-disrupting SigVars (Fig 4h; Methods). Using available single-cell human brain gene expression data (*45*), we annotated all members of this network for their highest expressed cell type (Methods), and found that all but one of these genes were expressed most strongly in neurons (Fig. 4i), suggesting that this disrupted PSP-associated signaling network functions primarily within neurons, consistent with the observation of overall neuronal enrichment of heritable PSP risk variants from GWAS (*46*).

### Refinement of SigVar annotations for high confidence predictions of causal variants

We next annotated the functional variants identified through our screen – representing likely causal variants – into those that fell within high confidence promoters or enhancers in neurons, astrocytes, microglia or oligodendrocytes (annotations (*36*)), as well as those variants that strongly disrupted predicted TFBSs (union of two algorithms (*43, 47*); Methods; Table S1). Of 320 SigVars, 233 (73%) had at least one additional functional annotation: 200 (63%) significantly altered TF-binding, 88 (28%) fell within a promoter or enhancer in at least one brain cell type, and 55 (17%) were double annotated for TFBS disruption and an additional promoter/enhancer mark. These 55 SNPs, disrupting TFBSs within annotated promoters or enhancers, represent the highest confidence causal functional variants within ten distinct GWAS loci, including *BIN1*, *HLA-DRB1/5*, *PTK2B*, *CLU*, *ICQK/KNOP1*, 17q21.31/*MAPT*, 19q13.32/*APOE*, *ASAP1*, SPI1/CELF, and *CASS4*.

### Systematic characterization of complex haplotypes 17q21.31 and 19q13.32

The chromosome 17q21.31 locus is noteworthy for harboring the tau-encoding *MAPT* gene within a common 900 kb inversion-polymorphism (*16*), and is a major risk locus for PSP (H1 haplotype OR = 4-5) as well as AD, Parkinson’s disease (PD), and Corticobasal Degeneration (CBD) (*12*). 17q21.31 contains complex haplotypic sub-structural variation and extensive LD, hampering interrogation with traditional statistical genetics approaches. We leveraged the ability to functionally dissect this region, testing 3,482 variants within 17q21.31 in strong LD with lead SNPs from a PSP GWAS (*10*), comprising approximately ∼24% of the more than 14,000 common variants in the region. Of these, we identified a total of 194 SigVars, of which 111 were stringently replicated in both MPRA experiments. Of these, 20 variants were also double annotated for active chromatin features and TFBS disruption, making them very high confidence causal regulatory variants (Supplemental Table S1).

We next clustered these SigVars based on LD (BigLD algorithm)(*48*), which identified seven distinct LD clusters within four contiguous LD blocks (Fig. 5a), suggesting distinct loci within this region. The largest LD cluster includes 42 SigVars within *MAPT* itself (-5 kb upstream of TSS to 3’ UTR) highlighting its striking regulatory complexity (Fig. 5b-c). Of these, 23 were replicated SigVars of which 13 are variants within annotated enhancers also predicted to disrupt TFBSs (Fig. 5b). The *MAPT* promoter region overlaps a large CpG island and a number of repetitive transposon elements that may impact gene expression (*49*). This region has been previously characterized using serial deletion assays in a variety of cell types (*50–52*), which identified a core promoter beginning -300 bp from the TSS (*49*). In this study, the region -226/- 63 (assay ID = 1447) was the 11^th^ most transcriptionally active library element assayed overall, while -349/-186 (ID = 1446) had only modest expression (Supplemental Data), suggesting a more restricted core promoter starting at -186 upstream of the *MAPT* TSS (Fig. 5d). We also identified SigVars within the broader promoter region (-4364/+3292; Fig. 5c) including rs17770296, which falls in the distal promoter (-2612) and overlaps a MLT1I (ERVL-MALR family) transposable element. Another variant, rs76324150 (+1485), falls within neuronal H3K4me3 peaks and is predicted to disrupt binding of the TF *ZFX* (Fig. 5d).

**Figure 5:**
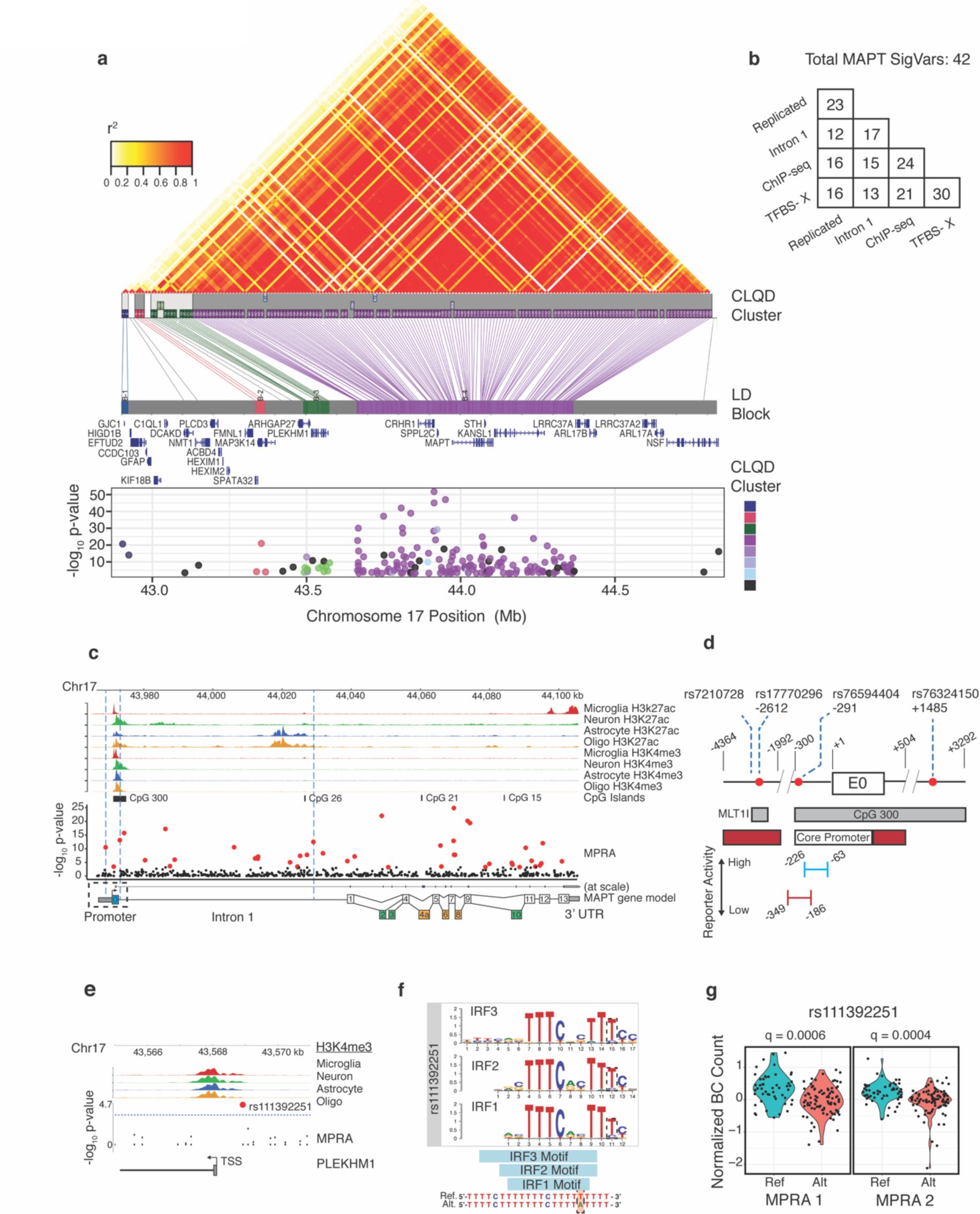
Systematic dissection of functional variation at 17q21.31. (**A**) Top: LD plot for common variants in 17q21.31 (1000 Genomes, CEU). MPRA SigVars were clustered by LD (CLQD clustering (*48*); Methods). Bottom: SigVars plotted by position across 17q21.31and MPRA significance (-log_10_ p-values; colors = cluster annotations, black variants are unclustered). Most variants fall within LD-cluster 4 (purple) centering on *MAPT*. (**B**) Annotation breakdown for *MAPT* SigVars. ChIP-seq from (*36*), TFBS-X = union of predicted TFBS-disruption from two algorithms (*43, 47*)). (**C**) Chromatin annotations and MPRA SigVars across *MAPT*. Genomic tracks (1–8) show H3K27ac (enhancer) and H3K4me3 (promoter) ChIP-seq peaks for microglia (red), neurons (green), astrocytes (blue), and oligodendrocytes (orange) from sorted human brain tissue (*36*), CpG islands (track 9), and all tested variants plotted by significance (- log_10_ p-value, track 10). SigVars (FDR q < 0.01, BH method) highlighted in red. Track 11 shows the MAPT gene model. Exons: Untranslated = blue, constitutive = white, alternative neuronal = green, rarely expressed in brain = yellow. (**D**) Functional annotations of the *MAPT* promoter (- 4364/+3292, numbering relative to TSS, features from (*52*)) with SigVars shown (red dots). Red = repressor regions. Bottom: MPRA relative reporter activity for two elements. (**E-G**) highlight rs111392251, a SigVar in the promoter (+/- 3kb) of *PLEKHM1*. (**E**) genomic tracks (1–4) for H3K4me3 peaks (as in **B**). (**F**) The alternate allele of rs111392251 is predicted to significantly disrupt the TFBS for *IRF1-3* TFs. (**G**) violin plots show normalized barcode distributions for each allele (MPRA FDR-adjusted q-values shown).

SP1 is known to bind the *MAPT* core promoter and regulate Tau expression (*53*), and our analysis also demonstrates that a SP1 signaling network contributes to PSP pathogenesis (Fig. 4g-i). Unfortunately, four potentially interesting variants within the proximal promoter region (- 144/+1485) dropped out of our assay likely due to the high GC content of the region (*49*) (Supplementary Text). Nevertheless, we identified four SigVars within the *MAPT* gene region predicted to disrupt binding of SP1 (Table S1), including rs76839282 which lies within H3K27ac peaks within the long regulatory intron 1 of *MAPT* (Fig. 5c).

We also identified SigVars within other LD blocks that are predicted to regulate independent risk genes. We highlight rs111392251, a high-confidence regulatory variant located in the promoter of *PLEKHM1* that is predicted to disrupt binding of *IRF-*family TFs (Fig. 5e-g)*. PLEKHM1* regulates autophagosome-lysosome formation (*54*) and has been previously suggested as a PSP risk gene (*9*), but has yet to be extensively characterized in a disease context. Other independent risk genes in distinct LD blocks implicated here include *MAP3K14* and *LRRC37A4P* (Table S2).

The *APOE* locus on 19q13.32 harbors the strongest common genetic association with late onset Alzheimer disease (LOAD), tagging the well-characterized *APOE4* risk haplotype (*6, 19, 20*). However, the extensive LD in the region coupled with the strength of the association signal has resulted in identification of hundreds of additional disease-associated variants (*18, 19*). Recent work involving transethnic scans and haplotype-aware conditional analyses have uncovered evidence for *APOE*-independent risk in the locus, implicating *PVRL2* and *APOC1* (*18, 55*), though others have argued that *APOE* coding variants mediate the entire association signal in the locus (*19, 20, 56*). We reasoned that identification of functional variants at 19q13.32 may help shed light on this complex regulatory architecture. We tested 640 variants in LD with the 538 genome-wide significant SNPs (*6*) at this locus, and identified 37 SigVars. Of these, at least 10 were whole blood or brain eQTLs for *PVRL2/NECTIN2* (GTEx)(*57*). These loci contained three intronic SigVars (rs34278513, rs412776, rs12972156), which were previously found to be significantly associated with LOAD when analysis was conditioned on *APOE4*-status (*18*). Our data identifies these variants as causal at this locus, and nominates their target gene as *PVRL2*. We also find that rs141622900, previously associated with cholesterol efflux capacity (*58*), is a SigVar residing in an active microglial enhancer directly downstream of *APOC1*, providing further support for *APOC1* as an AD risk gene. Finally, we identified an intergenic variant (rs2927437) within a robustly supported multi-tissue enhancer closest to the *BCL3* gene (*59*).

### Validation of select MPRA SigVars

We next selected five high-confidence causal variants residing within clearly annotated regulatory regions for additional validation, two variants within well-characterized AD loci near *BIN1* and *MS4A6*, and three variants within less well-described loci near *HLA-DRB1/5*, *CASS4*, and *ECDHC3*. We used CRISPR-Cas9 genome editing to assay regulatory regions in their native genomic context by excising the enhancer elements containing these variants, then assaying downstream effects on gene expression using quantitative PCR (Methods). We identified the target genes for a given variant either when the variant was close to the gene within its cis-regulatory region, or those linked by chromatin interaction data based on Hi-C from a relevant cell-type (Methods).

We first assessed rs13025717, which is predicted to strongly disrupt binding of KLF4, resides ∼20 kb from the TSS of the AD risk gene *BIN1*, and overlaps microglial and monocyte DHS as well as H3K27ac, H3K4me1, and H3K4me3 peaks (Fig. 6a-c). We used two pairs of gRNAs to excise a 240 or 374 bp segment containing rs13025717 in the SH-SY5Y cell line, and found a significant reduction in *BIN1* expression (Fig. 6d). Another intergenic variant, rs636317, lies within the Chr11:59923508 AD GWAS locus (*MS4A* gene region) (*6*) and overlaps H3K27ac and H3K4me1 peaks as well as DHS in microglia and monocytes (Fig. 6e). This variant is predicted to disrupt *CTCF* binding and is a highly significant MPRA SigVar (Fig. 6f-g). Analysis of existing Hi-C data from THP-1 cells (*60*) reveals rs636317 physically looping with upstream distal gene *MS4A6A*, which although not the closest gene to this variant, is also an eQTL (*57*) and the most highly-expressed gene at this locus in monocytes. As predicted, removing rs636317 in THP-1 cells, a macrophage-microglia related cell line (*60*) led to a significant reduction in expression of *MS4A6A* compared with both controls validating the function of this locus in a native context (Fig. 6h).

**Figure 6:**
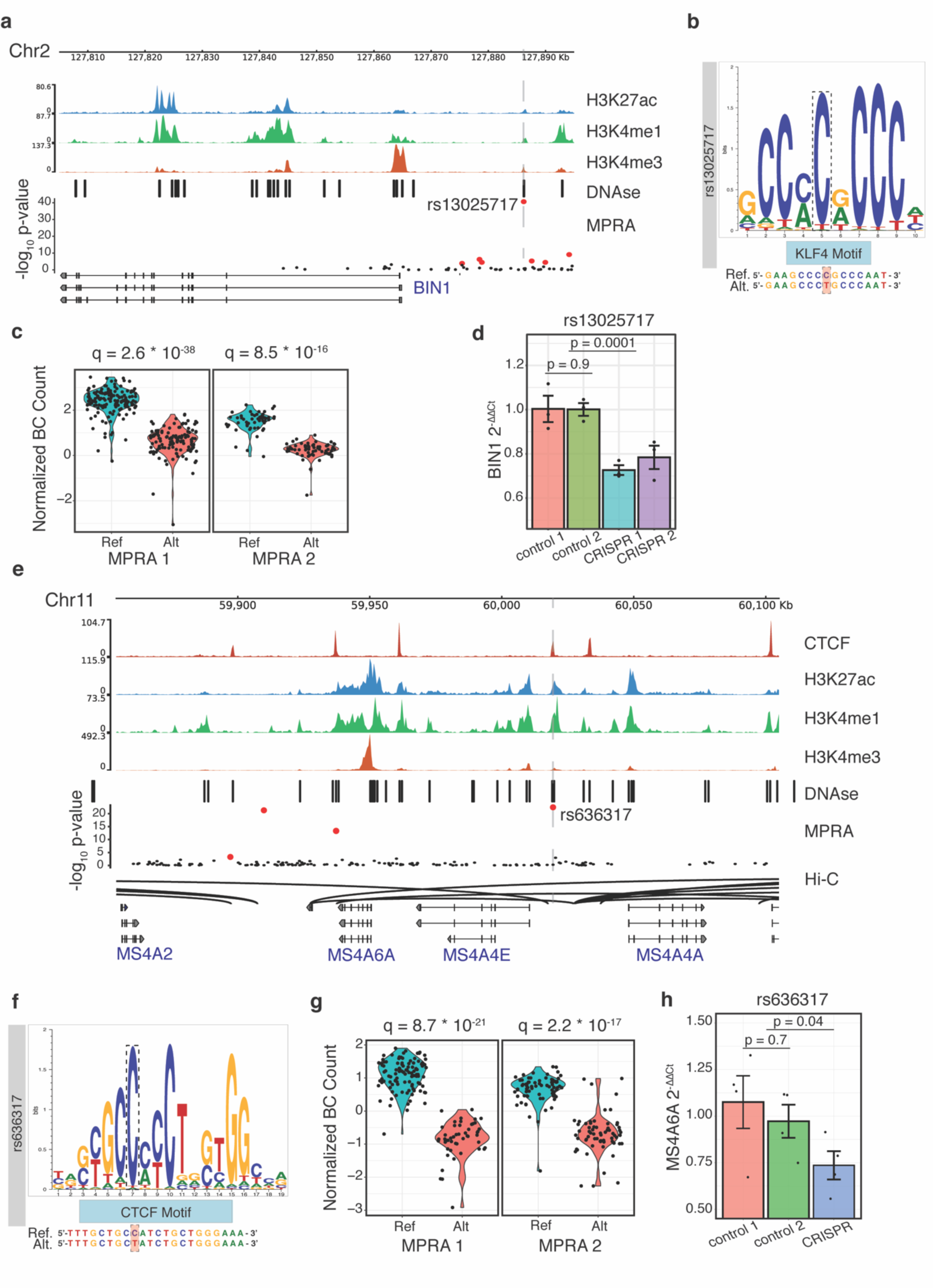
Validation of rs13025707 and rs636317 as regulatory variants underlying *BIN1* and *MS4A6* GWAS loci. (**A-D**) convergent evidence for rs13025707 as a regulatory variant. (**A**) Genomic tracks (1–4) at the *BIN1* locus show rs13025707 falling within H3K27ac, H3K4me1, H3K4me3, and DHS peaks (CD14+ monocytes; ENCODE). All variants tested by MPRA in the locus plotted by -log_10_ p-value (track 5). SigVars (FDR q < 0.01, BH method) shown in red. (**B**) the alternate allele of rs13025707 is predicted to disrupt binding of *KLF4*. (**C**) violin plots show normalized barcode distributions for each allele across both MPRA stages. FDR-adjusted p-values displayed. (**D**) CRISPR-mediated deletion of small genomic regions containing rs13025707 in SH-SY5Y cells significantly reduces *BIN1* expression compared with gRNA-scramble controls (n=3/group, combined n=6/condition; t(10) = -6.0; p = 0.0001; two-tailed Student’s t-test). Error bars = S.E.M. (**E-H**) convergent evidence for rs636317 as a regulatory variant. (**E**) Genomic tracks (1-5) at the *MS4A6* locus show rs636317 falling within CTCF, H3K27ac, H3K4me1, H3K4me3, and DHS peaks (CD14+ monocytes; ENCODE). All variants tested by MPRA in the locus plotted by -log_10_ p-value (track 6). SigVars shown in red. Monocyte Hi-C data (*60*) (track 6) links rs636317 with *MS4A6A*. (**F**) rs636317 is predicted to disrupt binding of *CTCF*. (**G**) violin plots show normalized barcode distributions for each allele across both MPRA stages. FDR-adjusted p-values shown. (**H**) CRISPR-mediated deletion of a genomic region containing rs636317 in differentiated THP-1 monocytes significantly reduces *MS4A6A* expression (n=4/group; t(10) = -2.3; p = 0.03; two-tailed Student’s t-test). Error bars = S.E.M.

The *CASS4* gene, although identified within an AD-risk locus for close to a decade (*6*), remains relatively understudied. We identified functional variant rs6064392 within a microglial enhancer in an intergenic region upstream of *CASS4* (significant *ATF-*family TFBS disruption; Fig. 7a-d). The minor T-allele of rs6064392 is in tight LD (r^2^ = 0.91) with the protective allele of rs6024870 (A; OR = 0.88; lead SNP)(*7*) and is predicted by MPRA to decrease downstream gene expression in agreement with eQTL data (Fig. 7c). We used a pair of gRNAs to excise a 382 bp region around rs6064392 in differentiated THP-1 cells, which significantly altered *CASS4* but not neighboring *RTF2* gene expression (Fig. 7e; Fig. S7a), supporting the functional impact of this variant on the *CASS4* gene.

**Figure 7:**
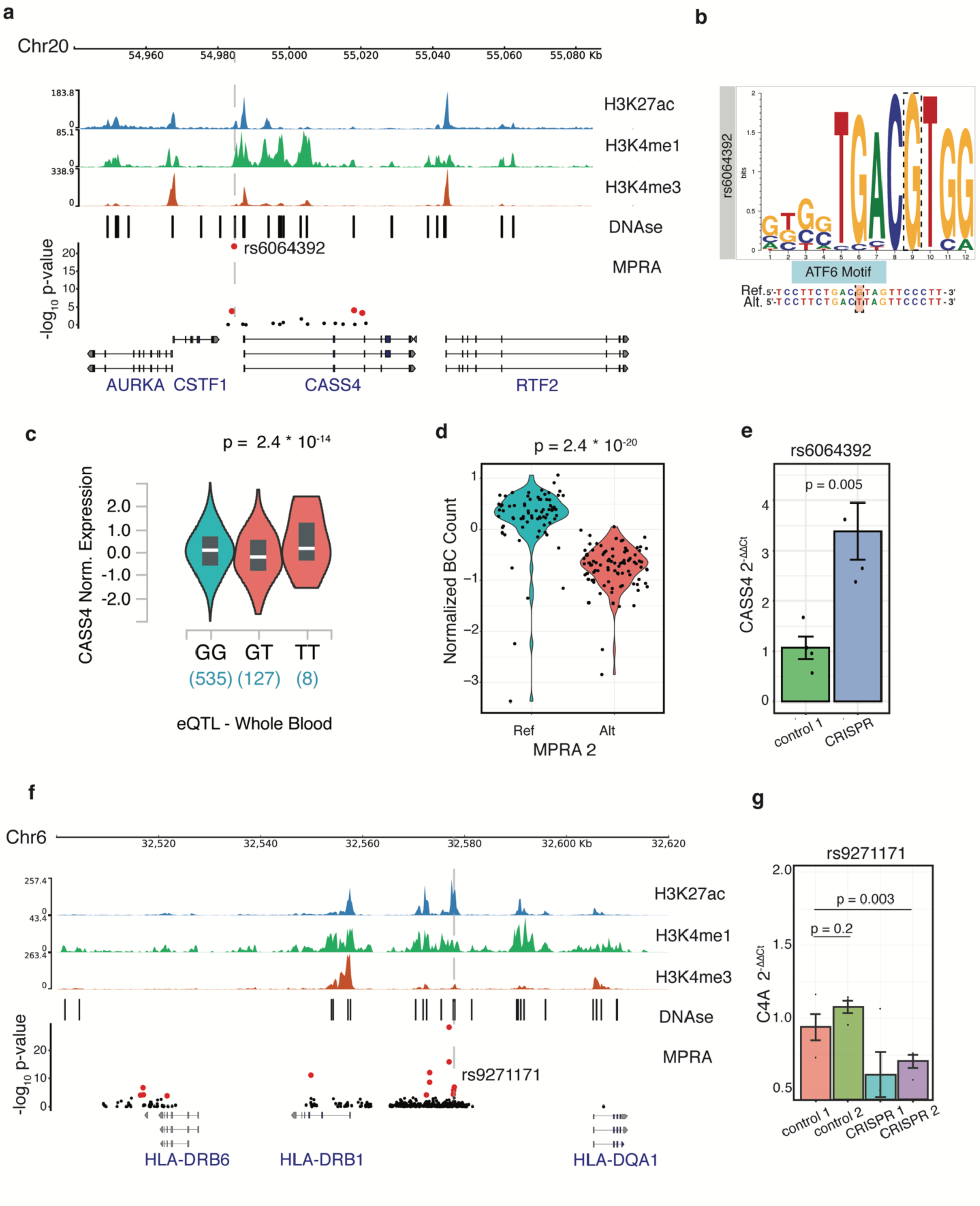
Validation of rs6064392 and rs9271171 as regulatory variants underlying *CASS4* and *HLA-DRB1/5* GWAS loci. (**A-E**) convergent evidence for rs6064392 as a regulatory variant. (**A**) Genomic tracks (1–4) at the *CASS4* locus show rs6064392 falling within H3K27ac, H3K4me1, and DHS peaks (CD14+ monocytes; ENCODE) upstream of the *CASS4* TSS. All variants tested by MPRA in the locus plotted by -log_10_ p-value (track 5). SigVars (FDR q < 0.01, BH method) shown in red. (**B**) rs6064392 is predicted to disrupt binding of *ATF6*. (**C**) Whole blood eQTL (GTEx) for rs6064392 and *CASS4*. (**D**) violin plot shows normalized barcode distributions for each allele (MPRA 2). FDR-adjusted p-values displayed (two-tailed Mann-Whitney-U test). (**E**) CRISPR-mediated deletion of a small genomic region containing rs6064392 in differentiated THP-1 cells significantly reduces *CASS4,* but not *RTF2* (Supplemental Fig 6a) expression compared with gRNA-scramble controls (n=4/group; t(6) = -4.3; p = 0.005; two-tailed Student’s t-test). Error bars = S.E.M. (**F**) Genomic tracks (1–4) at the *HLA-DRB1/5* locus show rs9271171 falling within H3K27ac, H3K4me1, H3K4me3, and DHS peaks (CD14+ monocytes; ENCODE). All variants tested by MPRA in the locus plotted by -log_10_ p-value (track 6). SigVars shown in red. (**G**) CRISPR-mediated deletion of genomic regions containing rs9271171 in differentiated THP-1 monocytes significantly reduces *C4A* expression relative to controls (n=4 group, n=8/condition; t(14) = 3.6; p = 0.003; two-tailed Student’s t-test). Error bars = S.E.M.

Next, we explored the *HLA-DRB1/5* AD GWAS locus, which is a highly polymorphic region of extended LD with numerous AD-associated variants and potential risk genes. Our screen identified nine significant variants in the locus, the majority of which fell within the intergenic region between *HLA-DRB1* and *HLA-DQA1.* We chose to further validate rs9271171 as it fell within a large myeloid open-chromatin region and was a significant eQTL in whole blood for multiple genes (GTEx)(*57*) where the alternate (protective) allele decreased reporter expression. We used two pairs of gRNAs to excise part of the enhancer region containing the variant in differentiated THP-1 macrophages, which revealed a dramatic (2.5-fold; Fig. S7b) increase in expression of *HLA-DQA1*. Interestingly, unpublished chromatin conformation data from PsychENCODE indicated that rs9271171 might also physically interact with the distal gene, complement 4 (*C4A),* for which it is also an eQTL in whole blood (GTEx)(*57*). Indeed, we observed that excision of this variant significantly reduced *C4A* expression in THP-1 macrophages (Fig. 7g). Its regulation of both *HLA-DQA1* and *C4A,* but in opposite directions suggested that this region may be pleiotropic and regulate multiple downstream genes. Because complement components are also expressed by reactive astrocytes, we performed the experiment again in human PSC-derived astrocyte cultures (Fig. S7c), confirming that excision of this variant reduced *C4A* gene expression in astrocytes as well. Lastly, the only variant that was tested, but not significant, rs7920721, was tested in THP-1 cells because of their similarity to microglia, but failed to impact gene expression for either *ECHDC3* or *USP6NL* (closest genes; Fig. S7d). However, we cannot rule out that this variant may function within a different cell type. In all, four of five variants significantly altered gene expression in the cis-regulatory region in the predicted gene of interest, validating these specific loci, and highlighting the general power of MPRA to identify functional variants.

## Discussion

Predicting functionality of noncoding variation is one of the major challenges in modern genetics. In this study, we provide the first systematic characterization of common variants underlying disease risk for two distinct neurodegenerative disorders: Alzheimer’s disease and Progressive Supranuclear Palsy. In doing so, we employ massively parallel reporter assays to screen 5,706 variants encompassing 34 unique loci identified across three genome-wide association studies (*6, 7, 10*), and obtain robust activity measures from 94% of tested elements. We find that active library elements are highly enriched for relevant chromatin and genomic features predictive of transcriptional regulation while confirming a high degree of inter- and intra-library reproducibility. We also utilize our two-stage study design to identify technical parameters impacting MPRA performance, adding to a growing literature on MPRA experimental design (*61, 62*) which will aid future studies (see Supplementary Text; Fig. S3c-d47:6; Fig. S8). Most saliently, we identify 320 variants with significant transcriptional skew between alleles at a conservative false discovery rate (q < 0.01), thus delineating putative causal variants at 27 of 34 tested genomic loci. We note that GWAS loci in regions harboring extensive LD and numerous risk genes are particularly difficult to interpret, and therefore require functional analyses. Here, we used MPRA to identify regulatory variants within two such regions, 17q21.31 and 19q13.32, risk factors for PSP and AD respectively, enabling identification of candidate regulatory variants. These data enabled us to identify regulatory variants within 29 and 17 distinct genes at 17qand 19q respectively (Table S3).

SNP-based heritability is known to be enriched within the regulatory regions within disease-relevant tissues (*63*). We demonstrate that functional regulatory variants further enrich within active, open chromatin, even relative to other GWAS variants within the LD block (Fig. 3e). Moreover, across two separate disorders, we find that a large proportion of functional variants fall within enhancers of distinct cell-types with a demonstrated relationship to disease, and that a large proportion of these enhancers are cell-type specific (71%; Fig. 3f-g).

We also determine that functional variants identified by MPRA are highly enriched for TFBS disruption, identifying transcriptional networks driving key aspects of disease risk (Fig. 4e-i; Fig. S6). For AD, this involves TFs including *NR4A2* and *SMAD-*family TFs, which have been described previously as acting within multiple cell-types in the brain and periphery to impact risk for AD (*64, 65*). For PSP, our analysis identified an enrichment of TFBS-disrupting functional variation in a significant protein interaction network with the transcription factor SP1. These TFs, as well as most of their predicted regulatory target genes are most highly expressed in neurons, consistent with cell-type specific aggregation of genetic risk in PSP. Moreover, we identified five SigVars in the PSP genome-wide suggestive locus harboring *SP1* (*10*), as well as four SP1 binding site-disrupting functional variants within the *MAPT* gene. These convergent findings provide strong evidence for disruption of SP1 signaling in neurons as a critical risk factor for PSP. SP1 is known to regulate a broad array of cellular processes including chromatin remodeling, apoptosis, immune regulation, and response to oxidative stress in neurons (*66*). While SP1 network dysregulation has been identified in AD brain (*67–69*) it does not harbor genetic risk, and therefore is likely to play a reactive or secondary role. Overall, our data are consistent with PSP risk primarily impacting neurons, and astrocytes and oligodendrocytes secondarily, and AD risk in microglia and astrocytes, as has been reported (*37, 46*).

Our observations also suggest a refined model for understanding common genetic risk. Signal from intergenic GWAS loci are typically interpreted as deriving from causal regulatory variants that influence downstream expression of specific cognate risk genes, and thus are explainable through colocalization with eQTLs. Our results, implicating a TF network converging on SP1 in PSP, are consistent with a model whereby common genetic variants function in aggregate across multiple TFBSs to disrupt key cell-type specific transcriptional programs. We speculate that this genetic mechanism may particularly manifest itself only upon induction of the relevant transcriptional network, which may occur within a disease context. For example, transcriptomic and proteomic studies have previously identified induction of an SP1 transcriptional network in tauopathies (*68, 69*), which here we show may interact with common variation at relevant binding sites distributed across the genome to determine risk. It has been previously observed that GWAS loci and eQTLs exhibit limited genetic sharing, leading to a “missing” mechanism of action explaining a large proportion of noncoding loci (*70*). Our data show that polymorphisms altering binding of critical, disease-relevant transcriptional networks offer an additional explanation. That these transcriptional networks regulate a large number of cell-type enriched genes, provides a mechanism whereby genetic risk is expressed, not by impacting a few core genes (*71*), but via polygenic cell type-specific regulatory effects on networks of genes (*72*).

A limitation of this study was the technical infeasibility of performing MPRA within human brain cell types, with the specific *trans*-regulatory environment of HEK293T cells likely influencing the generalizability of our screen (*29*). We addressed this by integrating cell-type specific regulatory annotations to prioritize variants likely to be functional within relevant brain cell types (Table S1). It should also be emphasized that this cell line showed the best overlap in active regulatory regions with multiple neuronal and glial cell types that contribute to disease risk, better than either a pure neuronal or glial cell type (Fig. 1b-c). We also identified and validated causal SNPs at four loci within brain-relevant cell lines, an important proof-of-principle for the external validity of our screen. In doing so, we provide strong evidence for rs13025717 and rs636317 as regulatory variants underlying the *BIN1* and *MS4A6A* loci respectively, as had been previously suggested (*73, 74*). Additionally, we identify rs6064392 as a likely causal variant within the *CASS4* locus, and our findings support previous reports that decreased *CASS4* expression may be protective in AD (*75*). Finally, we find evidence in two distinct cell types including IPSC-derived astrocytes that AD risk SNPs regulate *C4A*, the first time this gene has been implicated in common genetic risk for AD, further highlighting the importance of functional dissection of GWAS loci. We expect the combination of emerging eQTL and Hi-C data (*57, 76*) in conjunction with experimental CRISPR-mediated editing will enable further validation of other functional loci identified here, and their target genes. Additionally, some associated loci likely operate via mechanisms independent of direct transcriptional regulation, (e.g. splicing, post translational modification), requiring other approaches to delineate causal variants and their mechanisms. However, this study takes an important step forward by providing the first large-scale functional annotation of these dementia risk loci.

## Acknowledgements

We would like to thank Geschwind lab members Gayatri Nair and Samie Patel for their assistance with cell culture, and the UCLA TCGB and Flow Cytometry cores for their assistance and technical expertise.

## Author contributions

Conceptualization: YAC, GC, DHG

Data Curation: YAC

Formal Analysis: YAC

Funding Acquisition: YAC, GC, DHG

Investigation: YAC

Methodology: YAC, JED, SK

Project Administration: YAC, GC, DHG

Resources: SK, GC, DHG

Software: YAC

Supervision: SK, GC, DHG Validation: YAC

Visualization: YAC

Writing – original draft: YAC, DHG

Writing – review and editing: YAC, JED, SK, GC, DHG

## Data and materials availability

Raw and processed sequencing data for both MPRA experiments are available from GEO under accession GSE163855. To facilitate downstream analyses, preprocessed MPRA count data has also been uploaded to github: https://github.com/ycooper27/Tauopathy-MPRA. Processed and normalized MPRA data for both experimental stages are provided in Supplementary Table 1. Data underlying Figures 2c, 2d, 4a, 4c, 4g, 5a, 6d, 6h, 7e, 7g, and Supplementary Figures 4, 5, 6, and 7a are provided in Source Data.

## Code availability

Custom python scripts used for pre-processing MPRA data have been uploaded to github: https://github.com/ycooper27/Tauopathy-MPRA. Custom R code used for data analysis and visualization are available upon request.

## Materials and Methods

### Identification of candidate variants

<u>MPRA Stage 1</u>: We selected all genome wide significant (p < 5*×*10^-8^) variants from an AD and PSP GWAS (*6*),(*10*). We then identified all variants with a MAF > 5% in LD (r^2^ > 0.8) with these variants in Europeans (CEU + FIN + GBR + IBS + TSI; 1000 Genomes Phase 3) using the LDProxy tool accessed through the LDlink API. We subsequently filtered out indels, multiallelic, and coding variants. Both alleles of each variant were centered in 162 bp of genomic context (hg19/37) using the biomaRt (2.44.0) and BSgenome (1.56.0) packages in R (4.0.0) to create oligos. We then removed oligos containing KpnI, MluI, SpeI, and XbaI restriction sites needed for library cloning, leaving 5,223 total variants. Finally, we appended 5’ (CTGAGTACTGTATGGGCGACGCGT) and 3’ (GGTACCGACAAAAGTGTCAACTGT) PCR adaptor sequences to each oligo and synthesized the library on an Agilent 15k 210-mer array.

<u>MPRA Stage 2</u>: We replicated a selection of 186 SigVars (FDR adjusted q < 0.01) identified in MPRA 1 and 140 negative control variants. For 212 variants we also created oligos: 1) in reverse complement orientation, 2) with the variant located in the bottom third of the genomic context, and 3) in the reverse orientation (Supplemental Fig. 3c; Supplementary Text). Furthermore, in MPRA 2 we attempted to re-assess variants that dropped out of MPRA 1 (defined below). Finally, we assessed the lead SNPs from additional significant loci from two AD GWAS (*6, 7*), as well as 4 PSP genome-wide suggestive loci (*10*) (Table 1). LD partners were identified as above, constituting an additional 483 variants. The final MPRA 2 library was synthesized in duplicate on an Agilent 7.5k 210-mer array. All tested loci and oligo sequences for both MPRA stages are listed in Supplemental Information – Methods.

### Custom MPRA vector design

The pAAV-stop-MCS-bGH plasmid was created as follows: The multiple cloning site (MCS) and bGH-polyA sequence from the Donor_eGP2AP_RC plasmid (Addgene #133784) was cloned into the pAAV.CMV.PI.EGFP.WPRE (Addgene #105530) backbone using NheI and SphI restriction sites. To prevent transcriptional readthrough from the AAV ITR, this vector was re-cut at the NheI site and the transcriptional insulator element from pGL4.23 (Promega, E8411) was inserted using Gibson Assembly (*78*). The pMPRAdonor2-eGFP plasmid was created by cloning the eGFP open reading frame into the pMPRAdonor2 plasmid (Addgene #49353) digested with NcoI and XbaI restriction enzymes. Full plasmid sequences are provided in Supplemental Information - Methods.

### Library Construction

Sequences for primers described below are listed in Supplemental Information – Methods.

<u>Step 1:</u> We amplified and attached 20 bp degenerate barcodes to the oligo library by emulsion PCR (Chimerx, 3600-01). We performed four 50 uL PCR reactions with individual mixtures containing: 2 pmol of library, 1 uM of barcode_new_F primer, 1 uM of barcode_N_R primer, 200 uM dNTPs, 0.25 mg/mL acetyl-BSA (Thermo Fisher Scientific, AM2614), and 2 U of Phusion Hot Start II DNA Polymerase (Thermo Fisher Scientific, F549S) in 1X HF buffer. Thermal cycle conditions were: initial denaturation for 1 min at 95°C, followed by 20 cycles of 10 sec at 95°C, 20 sec at 61°C, and 20 sec at 72°C (2.5°C/sec ramp rate), followed by a final extension for 5 min at 72°C. Emulsions were broken with butanol, pooled, and purified per manufacturer’s instructions on spin columns. The amplified library and the pAAV-stop-MCS-bGH plasmid were digested overnight using SpeI-HF and MluI-HF enzymes and purified using Streptavidin M-270 Dynabeads (Thermo Fisher Scientific, 65305) or gel purification respectively (28704, Qiagen). An 80 uL T7 ligation reaction (NEB, M0318S) containing 200 ng of digested plasmid and 37.7 ng of library was performed followed by cleanup and electroporation into DH5*α* electrocompetent cells (NEB, C2989K). Transformed bacteria were pooled, serially diluted, and plated overnight at 37°C. Colonies from the dilution plate containing the number of bacterial colonies approximating 50-fold library coverage were collected and grown, followed by Maxiprep library extraction (Thermo Fisher Scientific, K210016).

<u>Barcode mapping</u>: Sequencing was performed to create a lookup table mapping barcodes to oligos. Oligo-barcode sequences were amplified from 2 ng of plasmid using flanking PCR primers (BC_map_P5_Rev and BCmap_P7_For) that added P5 and P7 adaptor sequences. Amplicons were sequenced by the UCLA TCGB core using an Illumina NextSeq 550 system (PE 2x150 bp) using custom Read 1 (BCmap_R1Seq_Rev) and Read 2 (BCmap_R2Seq_For) sequencing primers. Reads were merged using the BBMerge tool (*79*) and barcodes filtered and assigned to oligos using a python script. Briefly, reads that did not perfectly match library oligos were discarded. Barcodes represented by fewer than three reads were dropped. Ambiguously mapped barcodes were then filtered as follows: We bootstrapped an empirical distribution of oligo Levenshtein distances (python-Levenshtein 0.12.0) to determine a cutoff score (1^st^ percentile of distances). We then discarded barcodes where any pairwise read distance was greater than this cutoff score. The MPRA_barcode_mapping.py python script is provided: https://github.com/ycooper27/Tauopathy-MPRA.

<u>Step 2:</u> The MinP-eGFP fragment was amplified from the pMPRAdonor2-eGFP vector using Amp_minPLuc2_For and Amp_minPLuc2_Rev primers and both the plasmid library and fragment were sequentially digested using KpnI-HF and XbaI enzymes. These were used in a T7 ligation reaction containing 200 ng plasmid and 125 ng fragment. The ligation product was transformed into DH5*α* competent cells followed by plasmid isolation. The final library was configured with the 162 bp oligo upstream of the minimal promoter and the 20 bp barcode located in the 3’ UTR of the eGFP transcript (Fig. S1).

### Massively Parallel Reporter Assay

MPRA was performed with 6 biological replicates each consisting of ∼8 million HEK293T cells. We transfected 10 ug of library plasmid per replicate using Lipofectamine 3000 (Thermo Fisher Scientific, L3000008). 24 hours post-transfection, cells were dissociated, pelleted, and washed with DPBS. Replicates were lysed in 1 mL of RLT buffer and homogenized using QIAshredder columns (Qiagen, 79654). Total RNA was extracted and eluted in 100 uL RNase-free water using an RNeasy Mini Kit (Qiagen, 74104) with on-column DNase I digestion (Qiagen, 79254). We then extracted mRNA from 75 ug total RNA per sample using a Dynabeads mRNA Purification Kit (Thermo Fisher Scientific, 61006). Residual DNA was removed from 1 ug of mRNA using ezDNase (Thermo Fisher Scientific, 11766051) followed by reverse transcription using a custom primer (Lib_Hand_RT) and the SuperScript IV First Strand Synthesis Kit (Thermo Fisher Scientific, 18091050). RT was performed for 80 min at 52°C. We then amplified 10 uL of cDNA in 100 uL PCR reactions using NEBNext Ultra II Q5 Master Mix (NEB, M0544S) and Lib_Seq_eGFP_F2 and Lib_Hand primers for either 8 or 3 cycles (MPRA 1 and 2 respectively). Likewise, 5 replicates of 25 ng plasmid DNA (MPRA 1) or 4 replicates of 5 ng plasmid (MPRA 2) were amplified in 50 uL reactions using Lib_Seq_eGFP_F2 and Lib_Hand_RT primers. PCR products were purified using 0.6X-1.2X KAPA Pure Beads (Roche, KK8000) and then further amplified using P5_seq_eGFP_F2 and P7_Ind_#_Han primers for 7 PCR cycles to add P5, P7, and unique 8 bp Illumina index sequences. Following SPRI cleanup, amplicons were sequenced using an Illumina NextSeq 550 (1x20) or NovaSeq 6000 SP flow cell (SR 1x26 cycles) with 5% PhiX spike-in and custom Read 1 (Exp_eGFP_Seq_F2) and Index (Exp_Ind_Seq_P) primers.

### MPRA data analysis

#### Preprocessing

Raw reads were trimmed to only contain the 20 bp barcode and aligned to the previously generated oligo-barcode lookup table to create a barcode count matrix. Barcode reads that did not perfectly match were discarded. The MPRA_BC_counter.py mapping script is provided: https://github.com/ycooper27/Tauopathy-MPRA.

Barcodes were filtered such that at least 1 count had to be detected in every RNA replicate and at least 5 counts detected in each DNA replicate (to ensure stability of the log-ratios). A pseudocount of 1 was added to each barcode count, which was then normalized to sequencing depth to create counts per million reads per replicate (CPM). We then computed log_2_ RNA/DNA ratios for each barcode (DNA = median count across plasmid replicates), which were then quantile normalized between replicates using the preprocessCore (1.50.0) package. Variants with fewer than 5 unique barcodes for either allele were removed from further analysis.

#### Oligo activity measurements

To calculate transcriptional activity scores for each 162 bp oligo, allele-level summary statistics were computed as the median of the log_2_ barcode ratios. This value was then averaged between reference and alternate alleles to create an activity score for each oligo. To determine significance, this activity score was compared to the median activity value for the entire library using a one-sample Mann-Whitney-U test (two-tailed; n = 6 replicates), which was subsequently adjusted for multiple comparisons. “Active” elements for subsequent analyses had an increased RNA/DNA ratio compared with the library median at a Bonferonni adjusted p < 0.05 (or FDR adjusted q < 0.01 where described; Benjamini Hochberg method).

#### SigVar calculations

To identify variants with significant allelic skew (SigVars), log_2_ barcode ratios were combined across all 6 replicates by taking the median value. For each variant, a two-way Mann-Whitney-U test comparing barcode counts between each allele was used to identify allelic skew. SigVars were defined at an FDR threshold q < 0.01 (FDR adjustment, Benjamini-Hochberg method; further discussed below in Statistical Reporting). MPRA log_2_ effect sizes were defined as the median summed normalized barcode count for the alternate allele - reference allele.

#### Quality Control

Intra-experiment barcode reproducibility was defined as the mean pairwise correlation of each normalized barcode (RNA/DNA) count across all technical replicates. Allele correlation was determined by first finding the median normalized barcode count for each allele followed by determining mean pairwise correlation across all technical replicates (Both Pearson’s r and Spearman’s rho computed). Between experiment correlations for reference allele activity scores and variant effect sizes were also determined for 326 variants replicated in MPRA 2 estimate inter-experiment reproducibility.

#### Variant Dropout

Variants were excluded from analysis if we were unable to obtain activity measurements from at least 5 unique barcodes for both alleles (Fig. S8a; Supplementary Text). Of the 5,706 unique SNPs tested in both MPRA stages, 366 did not meet this inclusion threshold in either stage. We then compared sequence features from these “dropout” vs the rest of the “included” oligos. We used the SeqComplex perl module to compute GC content, CpG skew, and sequence complexity metrics (https://github.com/caballero/SeqComplex) (*80*). Sequence entropy was calculated using the RNAfold tool from the ViennaRNA 2.0 software suite (*81*). We also generated a custom python script to identify the longest runs of mono- or di-nucleotide repeats within each oligo (script provided: https://github.com/ycooper27/Tauopathy-MPRA). Outputted scores were compared between “dropouts” and all “tested” oligos using a two-sided Mann-Whitney-U test. All sequence scores are provided in Source Data. For visualization (Fig. S8) we took a random sample of 366 “tested” variants.

#### Power Analysis

We performed a power analysis to determine the sensitivity of our assay at different barcode complexities (Fig. S8b). We first determined empirical SigVar effect sizes from our combined study, which were binned into percentiles. We also determined an empirical assay standard deviation by taking the average standard deviation of the normalized barcode counts for all alleles that passed filter in both studies. We then performed a power analysis using the power.t.test function from the stats package in R, using the empirical standard deviation, an alpha threshold of p < 0.01and a “two.sided” hypothesis test. The analysis was performed using different percentiles (0, 20, 40, 60, 80th) of empirically computed effect sizes, as well as all integer n (i.e. # unique barcodes per allele) between 5-100, and plotted using ggplot2.

### Bioinformatic Analyses

#### Jaccard Index Calculations

We download DNase I hotspot files from the ENCODE project server (https://www.encodeproject.org/) (*82*) for brain cell types and tissues (cell types, accessions listed in Supplemental Information – Methods). Pairwise Jaccard indices between all samples were calculated using the Jaccard tool from BEDTools (2.29.2) (*83*) and plotted as a heatmap (Fig. 1b). Biological replicates (rep #2) were then discarded to avoid artificial inflation (leaving only one sample per cell type), and the average pairwise Jaccard index for each cell type with all other cell types was computed. GWAS overlap (Fig. 1c) was calculated for a given cell type by first intersecting all tested GWAS variants with that cell type’s DNase I peaks. Then, only the DNase-overlapping variants were intersected with DNase I peaks from the other cell types (pairwise) and the mean proportion shared was computed.

#### Chromatin annotation enrichment

We downloaded narrowPeak files for HEK293 DNase-seq, histone ChIP-Seq, and TF-ChIP-seq marks (accessions listed in Supplemental Information – Methods) from the ENCODE project server (*82*). We then determined overlap between these marks and MPRA “active” and “repressive” elements using the GenomicRanges R package (1.40.0) assuming a minimum of 1 bp overlap between the 162 bp oligo and the chromatin mark. Enrichment was calculated for active or repressive elements against a background set of all other tested oligos using a Fisher’s exact test, with log_2_ odds ratios and 95% confidence intervals reported. Active and repressive scores are provided in Source Data.

#### TFBS analysis

Predicted TFBS enrichment for active elements was calculated using the HOMER software suite (4.11) against a background set of all other oligos, after prefiltering for oligos containing reference alleles (Fig. 2e). In Fig. 4e, SigVar TFBS disruption was scored for all variants using the SNPS2TFBS webtool (*43*) and an enrichment odds ratio for TFBS-disrupting variants was calculated using Fisher’s exact test. For SigVars predicted to disrupt TFBSs, we correlated allelic skew effect sizes from our MPRA with predicted TFBS disruption scores. Sites with a TFBS score of 0 (denoting poorly defined scores) were discarded and the score with the max absolute value was chosen for sites with multiple predicted disruptions. We performed a similar analysis for SNPs from another large MPRA dataset (Fig. 4f) (*28*). As this dataset characterized variants embedded in genomic context from both positive and negative strands, we used the max observed effect size. We also partitioned SigVars into those that were derived from AD or PSP GWAS (111 or 209 variants respectively; 17q21.31 variants were considered PSP), and re-ran them through the SNPS2TFBS algorithm (Source data). The output was filtered for TFBSs with at least two predicted disruptions and enrichment p-values were then FDR-adjusted (BH method). Only TFBSs significantly (q < 0.05 or q < 0.1) enriched for disruption by SigVars were plotted using ggplot2, with disruption counts, log_2_ enrichment odds ratios, and unadjusted -log_10_ p-values shown (Fig. 4g; Fig. S6).

We also tested whether HEK293T cell TF abundance correlates with MPRA allelic skew. To do so, we scored all variants tested in MPRA 1 using the SNPS2TFBS software. For each variant predicted to disrupt a TFBS, we found the corresponding TF’s normalized gene expression in HEK293T cells using RNA-seq data from the human protein atlas (*84*) (accession in Supplemental Information – Methods). We then found the Spearman’s correlation between TF expression and the MPRA absolute value log_2_ fold change.

For the 6 TFs significantly disrupted in PSP (at q < 0.05) or 14 TFs at q < 0.1, we created a protein-protein interaction network using the STRING (v11) webtool and standard parameters (Fig. 4g; Fig. S6) (*77*). We then computed empirical SP1-connectivity p-values for this network using resampling as follows: We determined protein-protein interactions for all TFs annotated by the SNPS2TFBS tool (165 TFs total; output in Source Data – Fig 4g.3) using STRING (https://string-db.org). We then found the number of edges between SP1 and either 5 or 13 additional randomly sampled TFs, repeating this procedure 10,000 times to create a distribution, which was compared to the true number of PSP-network edges (4 or 8 respectively) to generate a SP1-connectivity p-value. We then identified all sites among the 209 PSP-SigVars predicted to disrupt binding of these 6 TFs (union of SNPS2TFBS and motifbreakR annotations; Supplemental Table 1) and found all genes within +/- 10 kb of these sites, which we called “target genes” (Fig. 4h). We then annotated the network composed of these TFs and their targets using single cell RNA-seq data from human M1 cortex (© 2010 Allen Institute for Brain Science. Allen Human Brain Atlas. Available from: https://portal.brain-map.org/atlases-and-data/rnaseq/human-m1-10x) (*45*) . We annotated each network gene as belonging to the corresponding cell type with the highest trimmed mean gene expression value, and created a boxplot displaying the number of annotated genes per cell type (Fig. 4i).

We also tested whether TFBS disruption was predictive of MPRA allelic skew. For this analysis, we considered variants derived from the external MPRA dataset (*28*). Only variants with computed allelic skew values were considered. We used the maximum -log_10_ p-value for variants tested in multiple configurations and also filtered out variants missing valid rsIDs. We then performed FDR adjustment (BH method) on the MPRA-determined allelic skew p-values, and labeled SigVars at q < 0.01 thresholds. All variants were scored for TFBS disruption using the motifbreakR package (2.2.0) utilizing the HOCOMOCO v10 TF binding model (*47, 85*) (“strong” effect, and binding threshold of p < 1*×*10^-4^). Variants that could not be identified in dbSNP v144 were not scored and were subsequently discarded. We then computed the positive predictive value between predicted TFBS disruption and SigVar labels. We also downloaded RNA-seq data for K562 cells from the human protein atlas (*84*), and identified the top 200 TFs (by normalized counts) expressed in these cells. We then filtered for “strong” predicted TFBS disruption only for motifs corresponding to these top expressed TFs, and then re-computed PPD.

Finally, we tested whether disruption of particular TFBSs predicted MPRA allelic skew (Fig. S5). We again only considered K562 expressing TFs. For each collection of variants predicted to disrupt a particular TFBS, we found the proportion that were also annotated as SigVars (SigVars defined at q < 0.01). We then assessed the statistical significance of these proportions for each TF by performing a one-tailed binomial test against the background SigVar probability in the overall dataset (prob = 0.115), leaving a p-value which was FDR adjusted. Enrichments are shown in Source Data.

#### Comparison with computational predication algorithms

We scored all tested variants using the LINSIGHT, CADD, CATO, and GWAVA algorithms (*39–42*), using each algorithm’s precomputed scores. When an algorithm provided multiple scores per variant (particularly LINSIGHT), scores were averaged. Individual variant scores are provided in Source Data. First, we correlated (Spearman’s rho) MPRA effect-sizes with computational predicted scores. We then labeled MPRA SigVars at FDR-adjusted q < 0.01 thresholds and calculated AUC for each algorithm using the ModelMetrics (1.2.2.2) package. We then performed binary classification on all scored variants by positively labeling MPRA SigVars (q < 0.01) and an equivalent number of the top scoring variants from each algorithm before calculating pairwise Cohen’s Kappa using the ModelMetrics package. These analyses were replicated using another published MPRA dataset (*28*) as above.

#### SigVar functional annotations

SigVars from this study were annotated for TFBS disruption and overlapped with functional brain annotations (Supplemental Table 1). We calculated TFBS disruption using both the motifbreakR package using the HOCOMOCO v10 TF binding model (filtered for a binding threshold of p < 1*×*10^-4^ and “strong” predicted effects) as well as the SNPS2TFBS webtool (*43*). Additionally, published enhancer and promoter annotations for sorted microglia, neurons, astrocytes, and oligodendrocytes were downloaded and converted to bed files using the ucsc-bigbedtobed tool and overlapped with SigVars using BEDTools intersect (*36, 83*). We also identified SigVars overlapping high-confidence multi-tissue enhancers defined by the HACER database (*59*). Promoter/Enhancer accessions are provided in Supplemental Information – Methods.

#### LD clustering

To calculate LD between SigVars within the 17q21.31 locus, we downloaded chr17 VCF files from the 1000 Genomes FTP server. The VCF was subsetted for 90 unrelated individuals of CEU ancestry and reformatted to get cumulative allele frequencies using PLINK 1.9 (*86*). Genotype files were loaded into R and used to create LD clusters for SigVars within 17q21.31 using the clqd function (CLQDmode = “maximal”) from the gpart R package (1.6.0) (*48*).

### Cell Culture

We obtained HEK293T (CRL-3216), THP-1 (TIB-202), and SH-SY5Y (CRL-2266) cell lines from ATCC. HEK293T cells were cultured in DMEM containing GlutaMAX (Thermo Fisher Scientific, 10566016) supplemented with 10% FBS and 1% Sodium Pyruvate (11360070). THP-1 cells were cultured in RPMI-1640 medium (ATCC 30-2001) supplemented with 10% heat-inactivated FCS and 10 mM HEPES buffer. THP-1 differentiation was performed through addition of 20 ng/mL of PMA (Millipore Sigma, P1585) to the culture media for 48 hours. SH-SY5Y cells were cultured in 1:1 EMEM/Ham’s F12 (ATCC 30-2003 / Thermo Fisher Scientific, 31765035) supplemented with 15% HI-FCS, 1% Sodium Pyruvate, and 1% MEM-NEAA (Thermo Fisher Scientific, 11140050). For differentiation, SH-SY5Y cells were gently passaged and plated on Poly-D Lysine coated plates. After 24 hours culture media was replaced with a differentiation media composed of Neurobasal A (Thermo Fisher Scientific, 10888022), GlutaMAX, B27 Supplement (17504044), and 10 uM Retinoic Acid (Millipore Sigma, R2625). Cells were differentiated for 5 days with half-media replacement every 48 hours.

Induced Pluripotent Stem Cells (IPSCs) used in this study were previously documented (*87*) and kindly provided by Dr. Li Gan in accordance with the UCLA TDG guidelines. IPSCs were differentiated into mature astrocytes for 120 days as previously described (*88*), and were maintained post-differentiation in DMEM supplemented with 1% Sodium Pyruvate, 10% HI-FCS, and 1x N2 supplement (Thermo Fisher Scientific, 17502048) until use.

### CRISPR experiments

We excised enhancers containing rs636317, rs13025717, rs6064392, rs9271171, and rs7920721 as follows: Pairs of guide RNAs targeting upstream (5’) and downstream (3’) flanking sequences were designed and cloned into LentiCRISPRv2-GFP (Addgene #82416) and LentiCRISPRv2-mCherry (#99154) respectively using the BbsI restriction site (gRNA sequences in Supplemental Information – Methods). Lentiviral particles were produced in HEK293FT cells by triple transfection as previously described (https://www.addgene.org/protocols/lentivirus-production), concentrated using Lenti-X (Takara Bio, 631232), and resuspended in DPBS. Guide pairs were then screened for cutting efficiency resulting in two pairs targeting rs13025717 (gRNA #s: 11 + 12, 11 + 14), one pair targeting rs636317 (15 + 16), one pair targeting rs6064392 (25 + 26), three pairs targeting rs9271171 (3 + 4, 5 + 4, 5 + 6), and one pair targeting rs7920721 (21 + 20). Guide pairs or scramble gRNA control lentiviruses (MOI ∼ 0.5) were then used to infect 80% confluent 6-well plates of SH-SY5Y cells or t25 flasks of THP-1 cells. Culture media was replaced 16 hours later and cells were expanded for 5 days post-infection. Cells were sorted at the UCLA BSCRC flow cytometry core to isolate ∼500,000 GFP+/mCherry+ cells per replicate, and were subsequently differentiated. Ultimately we excised a 240 or 374 bp region containing rs13025717 in SH-SY5Y cells, a 430 bp region containing rs636317 in THP-1 cells, a 382 bp region containing rs6064392 in THP-1 cells, a 1065 or 682 bp region containing rs9271171 in THP-1 cells (as well as a 1005 bp region in IPSC-derived astrocytes), and a 473 bp region surrounding rs7920721 in THP-1 cells.

For each replicate we collected total RNA and genomic DNA using the AllPrep DNA/RNA Mini Kit (Qiagen, 80204). CRISPR-mediated removal of the target enhancer was assessed by amplifying the target region of gDNA by PCR (genomic PCR primers: Supplemental Information - Methods) and verifying strong representation of the truncated allele via gel electrophoresis. cDNA was reversed transcribed using SuperScript IV, Oligo(dT)_20_ primer, and 300ng of total RNA. We performed qPCR using the KAPA SYBR FAST Kit (Roche, KK4600), 500nM qPCR primers (qPCR primers: Supplemental Information - Methods) and a Roche LightCycler 480. Relative transcript abundance was quantified using the 2^-ΔΔCT^ method (*89*) normalized to the geometric mean of ACTB and GAPDH reference genes.

### Statistical Reporting

Statistical analysis was performed using the stats package in R. All hypothesis testing was two-sided except for the binomial test used in Supplemental Fig. S5. Unless otherwise stated, all enrichment analysis was performed using a Fisher’s exact test. MPRA allelic skew multiple testing correction was performed as follows: Variants from MPRA 1 were combined with additional unique variants tested in MPRA 2 (total 5340 variants) and Mann-Whitney-U p-values were FDR-adjusted (BH method). SigVars were called at a threshold of q < 0.01. To assess variant reproducibility, variants replicated*, i.e.* re-tested (320 total) in MPRA 2 were considered separately, and assigned significance at a Bonferroni-adjusted q < 0.05.

### Data Visualization

Variant genomic annotations were determined and plotted using the annotatr (1.14.0) Bioconductor package and ggplot2 from the tidyverse collection (1.3.0). Heatmaps were generated using the pheatmap R package (1.0.12). The circle Manhatten plot was created using the CMplot R package (https://github.com/YinLiLin/R-CMplot). ROC plots were visualized using the plotROC (2.2.1) extension and ggplot2 (*90*). 17q21.31 LD plots were created using the BigLD function from the gpart package (*48*). Genomic tracks and chromatin annotations were visualized using the pyGenomeTracks python module (*91*) . All other data were visualized using ggplot2 with the GGalley package extension.

## Supplementary Text

### Massively Parallel Reporter Assay Technical Considerations

In this supplemental note, we provide further discussion and analysis of factors observed to impact MPRA experimental outcomes and technical performance. Consideration of these parameters may aid in the design and implementation of future studies involving complex library construction and massively multiplexed assays.

### MPRA reproducibility

Because of our two-staged experimental design, we took the opportunity to characterize the true reproducibility of our assay. We identified 326 variants in the first MPRA stage, including 186 with significant transcriptional skew between alleles (SigVars; FDR adjusted q < 0.01) to be re-tested in the second stage. Importantly, although library design stayed identical, oligo barcoding, cell culture, sequencing, and other technical factors were distinct between stages, thereby providing an informative estimate of the impact of technical confounding variables. Fortunately, we found that our assay was highly reproducible, with measurements of allele transcriptional efficacy, and relatedly, MPRA-determined effect sizes, highly correlated between experiments (r = 0.98 and 0.94 respectively, both p < 2*×*10^-16^; Fig. 3a). This suggests that MPRA-determined measurements of transcriptional efficacy are highly precise and that variant ordering (e.g. prioritization) on the basis of effect size is robust. Significantly, reproducibility is not an inherent feature of SigVars or highly expressed variants in particular. Upon inspection of the 140 re-tested variants without significant allelic skew (q > 0.01), we again confirmed high correlation between stages (r = 0.96, p < 2*×*10^-16^)

Interestingly, 152 of 186 (82%) re-tested MPRA 1 SigVars remained significant in stage 2 (replication threshold; Bonferroni q < 0.05; Fig. 3a), which is somewhat lower than might be expected considering the original conservative FDR threshold of q < 0.01 used to determine significance in stage 1 and the near perfect correlation of effect sizes observed between experiments. In comparing the 34 non-replicating and 152 replicating variants, the non-replicates had a *higher* MPRA stage 1 median barcode complexity (least complex allele; 94 vs 67 barcodes), and a *lower* MPRA stage 2 barcode complexity (45 vs 65). Thus, variants that failed to replicate had a dramatic reduction in barcode complexity in the replication stage (delta barcodes = -49 vs -3 barcodes). Concurrently, non-replicating variants had a significantly lower mean absolute effect size (MPRA stage 1 Log_2_ FC: 0.36 vs 0.72, p = 2.6 *×*10^-9^; two-sided Mann-Whitney-U test). We next determined power to detect significant allelic skew for a range of percentiles of empirical MPRA effect sizes at different levels of barcode complexity (Fig. S8b; Methods). Interestingly, the threshold at which 80% power is achieved for the 40^th^ percentile of empirical effect sizes is at a barcode complexity of 42 barcodes per allele, with a steep drop-off in power at decreasing effect sizes and barcode complexities. Thus power to detect SigVars of small effects are highly contingent on high (>40 barcodes/allele) barcode complexity, and suggests that while MPRA is highly precise in determining allele effect sizes (even at low complexities ∼5 barcodes, Data not shown), identification of allelic skew *significance* will be somewhat impacted by stochastic fluctuations in barcode complexity due to PCR.

### Effects of oligo configuration on MPRA performance

We also tested the effects of library construction parameters on SigVar detection and reproducibility, using the SigVars identified in MPRA stage 1 as a “gold standard” test set. Specifically, we tested the impact of placing the oligo in the reverse complement (RC) orientation in the MPRA vector as well as the effect of placing the variant in the bottom third (as opposed to the middle) of the 162 bp genomic context. We used the oligo in reverse orientation as a negative control as this maintained identical nucleotide composition (Fig. S3c). Interestingly, SigVars had strong but imperfect correlation with their RC (r = 0.69, p < 2*×*10^-16^) and lower third (r = 0.78, p < 2*×*10^-16^) counterparts, suggesting modest effects of oligo orientation and distal sequence context on MPRA activity. As MPRA activity measures are highly precise and reproducible (discussed above), these effects of oligo configuration are likely biological and not due underlying assay noise. Placing oligos in the reverse orientation completely abolished activity as expected (all p > 0.5; Fig. S3c). These findings can inform design considerations for future massively parallel screens.

### Technical factors influencing variant drop-out

We initially assumed that the observed 9% variant drop-out during stage 1 was random bottlenecking due to library construction and therefore attempted to re-test missing variants in stage 2. However 346/491 of these variants also failed to pass QC in stage 2, suggesting that a large proportion of drop-outs were due to systematic amplification failure during library construction or strong transcriptional repression. Indeed, of the vast majority of variants that dropped out specifically during MPRA 1 library construction dropped out again during MPRA 2 library construction, suggesting PCR amplification failure. We assessed sequence features of the 366 unique variants that failed to pass quality control (QC) for both MPRA 1 and 2 and found that missing variants were more likely to contain GC content greater than 75% or less than 25% (GC extremes) and had increased rates of CpG skew. Additionally, drop-out variants had significantly lower mean sequence complexity (measured by Shannon’s entropy, compressibility, linguistic complexity measures, *etc.*; Methods), and were more likely to contain long regions of repetitive sequences such as CpGs, dinucleotide repeats, or polyA tracts (Fig. S8a).

### Library expression impacts assay performance

It has been previously reported that MPRA performance, as defined by the precision and reproducibility of measurements of library transcriptional efficacy, is dependent on high levels of library expression (*33*). This is particularly true for assays measuring variant function, which attempt to identify subtle changes in transcriptional efficacy between alleles (i.e. detect small effect size events). Three critical factors influencing expression of reporter libraries are: 1) the ability of library elements to drive transcription, 2) global transcriptional propensity of the particular cell-type, 3) transfection or infection efficiency into cells of interest (e.g. copy number per cell, cell transduction percentage). Unfortunately, assays testing variant allelic skew typically employ weak minimal promoters so as not to overwhelm the transcriptional impacts of single nucleotide substitutions, which are typically small. As a result, these assays have been exclusively performed in cancerous cell lines, which can be transfected at high efficiencies. Moreover, cancer cells, particularly those with c-Myc amplifications have been noted to have higher levels of overall transcription (known as transcriptional amplification) (*92*).

We confirmed the importance of library expression on assay performance by comparing four separate MPRA experiments: the two assays presented in this study as well as two unpublished assays. We took the average between-replicate correlation (Spearman’s rho) of allele expression as the overall performance metric for each experiment. Significantly, each experiment required a differing number of PCR cycles to amplify an adequate amount of library cDNA for downstream sequencing (10, 15, 17, 18 total PCR cycles). We found an inverse relationship between PCR cycle number and performance (mean rho; Fig. S8c), with a large performance drop off above 15 cycles. As PCR cycle number likely reflects underlying mRNA quantity and library expression, this confirms the importance of expression on performance. This observation represents a major technical obstacle towards implementing MPRAs within difficult to transduce primary cell-lines, which may be desirable *in vitro* model systems that more closely recapitulate relevant biology than cancer lines.

---

**Figure S1:**
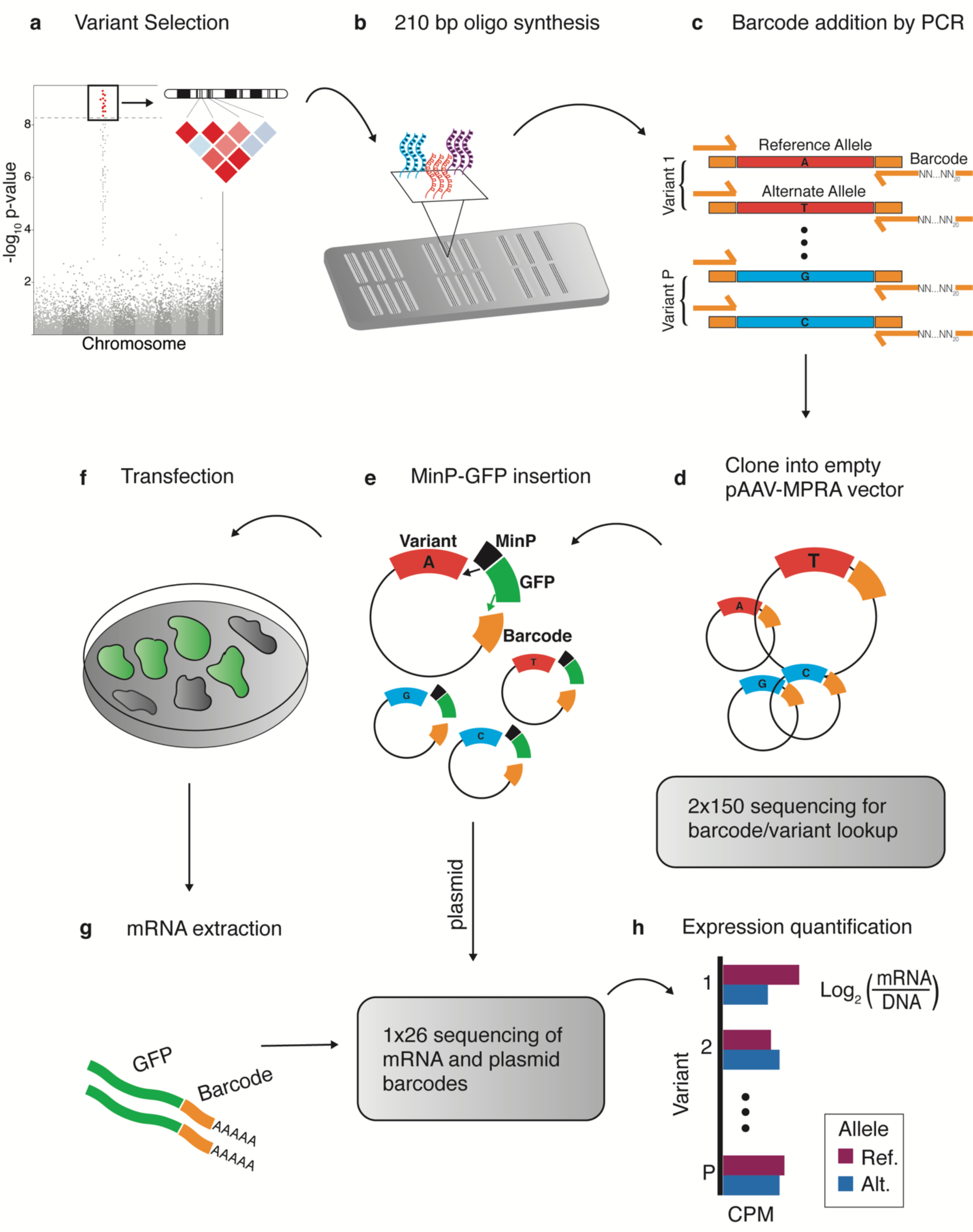
Technical schematic for massively parallel reporter assay (MPRA). (**A**) We selected all bi-allelic noncoding genome-wide significant (p > 5*×*10^-8^) or locus lead SNPs and LD partners (r^2^ > 0.8) from three GWAS (Methods). (**B**) The reference and alternate alleles of each variant were centered within 162 bp of genomic context (hg19) and synthesized on an Agilent array. (**C**) Oligos were amplified by PCR using primers complementary to flanking shared adaptor sequences. This step also attached 20 nucleotide random barcodes and restriction enzyme sites. (**D**) Oligos were digested and cloned into an expression library, followed by paired-end 2x150 bp sequencing to map barcodes to associated oligos. (**E**) A minimal promoter and eGFP gene was inserted between the oligo and barcode, with the oligo upstream of the minimal promoter and the barcode in the 3’UTR of eGFP. (**F**) The expression library was transfected into HEK293T cells for 24 hours. (**G**) following mRNA extraction, mRNA and plasmid barcodes were amplified, pooled, and sequenced using indexed single-end sequencing. (**H**) mRNA barcode counts were normalized to DNA counts, and the median normalized barcode count was taken as an activity summary score for each allele.

**Fig. S2:**
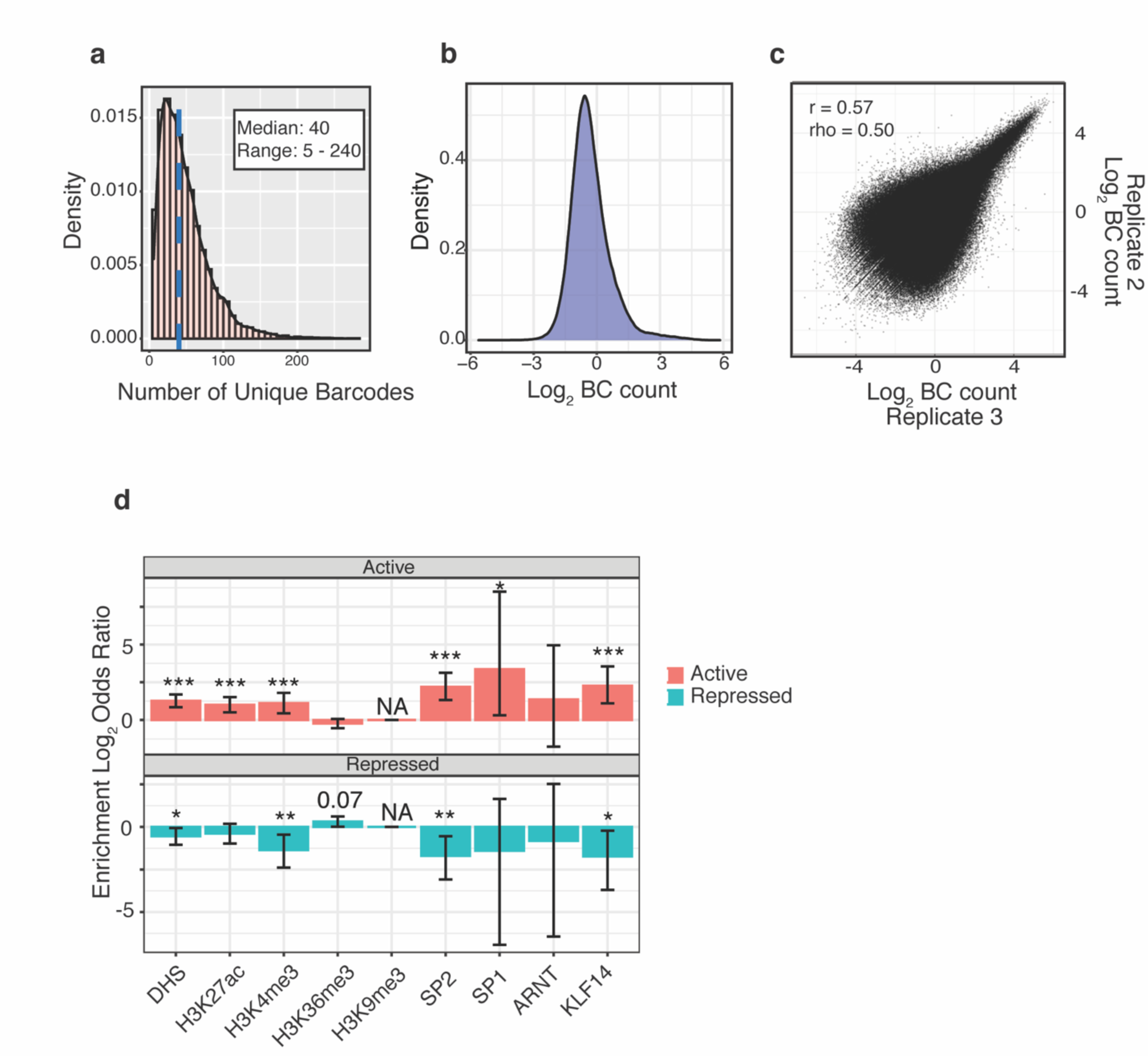
MPRA 1 extended quality control metrics. (**A**) Density plot shows the distribution of uniquely mapped 20 nucleotide barcodes per individual allele (i.e. unique barcodes/allele; 9,464 total alleles) for the full MPRA 1 library, blue line = median (40 barcodes/allele). (**B**) Density plot of log_2_ normalized barcode (BC) ratios. (**C**) Representative plot showing the correlation of log_2_ normalized barcode counts between technical replicates (Pearson’s r = 0.57; Spearman’s rho = 0.50; both p < 2*×*10^-16^). (**D**) Enrichment log_2_ odds ratios (Fisher’s exact test) of active and repressed elements (activity defined at FDR q< 0.01, BH method) within HEK293T ChIP-seq peaks for both histone and TF marks. Error bars = 95% CI, *** FDR-adjusted p < 0.001 (BH method), ** p <0.01, p < 0.05.

**Figure S3:**
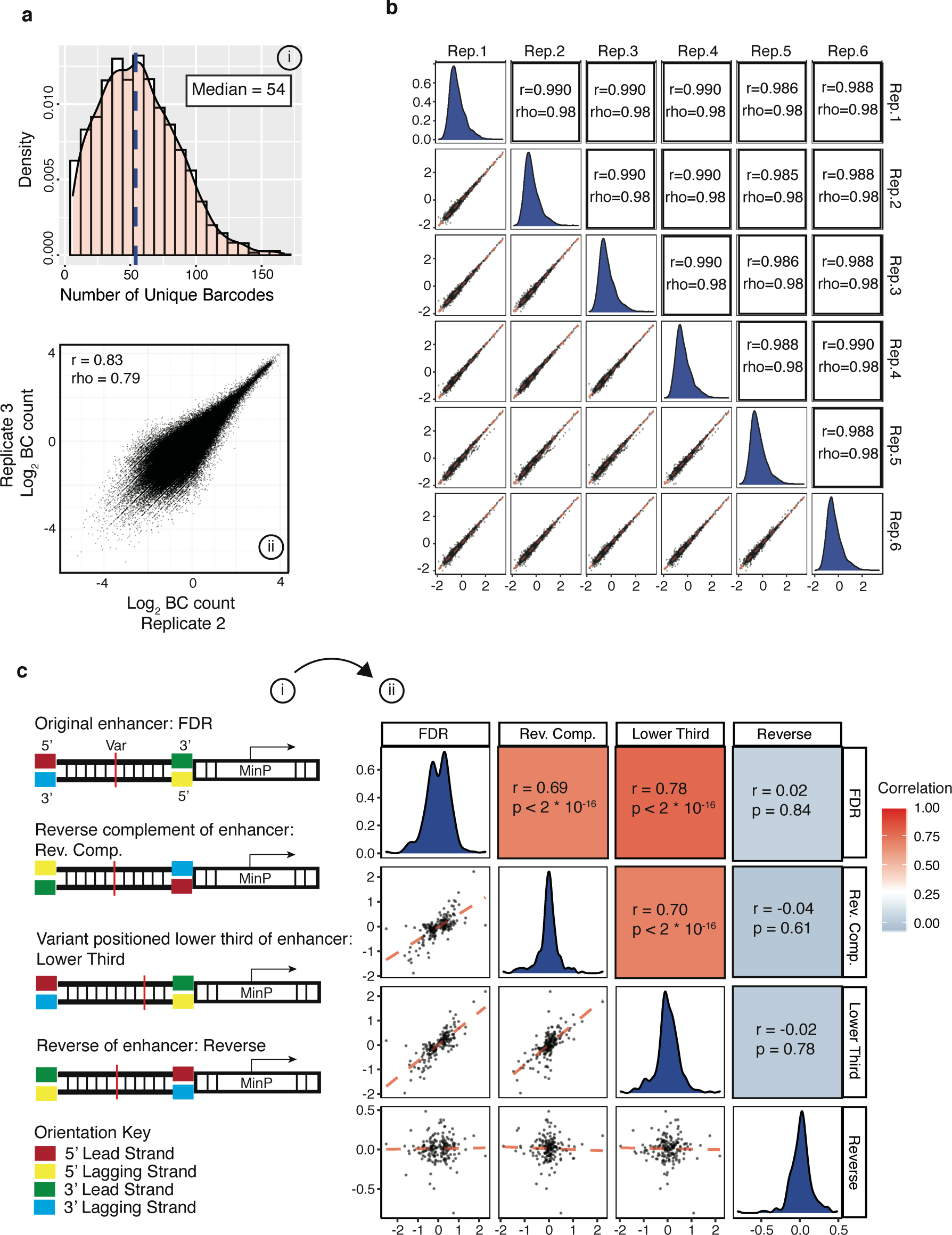
MPRA 2 quality metrics. (**A**) Barcode distribution for the 3,072 unique alleles tested in the full MPRA 2 library (blue line = median) with ii) representative plot showing correlation of normalized barcode (BC) counts between technical replicates (Pearson’s r = 0.83, Spearman’s rho = .79, p < 2*×*10^-16^). (**B**) Panels show pairwise correlation of median log_2_ normalized barcode counts for all alleles passing filter (n = 3,072) between 6 technical replicates. Red line = OLS regression line of best fit, all p < 2*×*10^-16^. (**C**) i) Shows the experimental design for testing the effect of enhancer orientation and variant placement on MPRA activity. 212 variants identified from MPRA 1 were replicated in MPRA 2: Oligos were kept in the same orientation as the original (FDR), oligos were placed in the reverse complement orientation (Rev. Comp.), variant was placed in the lower third of the oligo rather than the middle (Lower Third), or the enhancer sequence was reversed (Reverse) as a negative control. Plots showing correlations between oligo orientations shown in ii), red line = OLS regression line of best fit, Pearson’s correlation.

**Figure S4:**
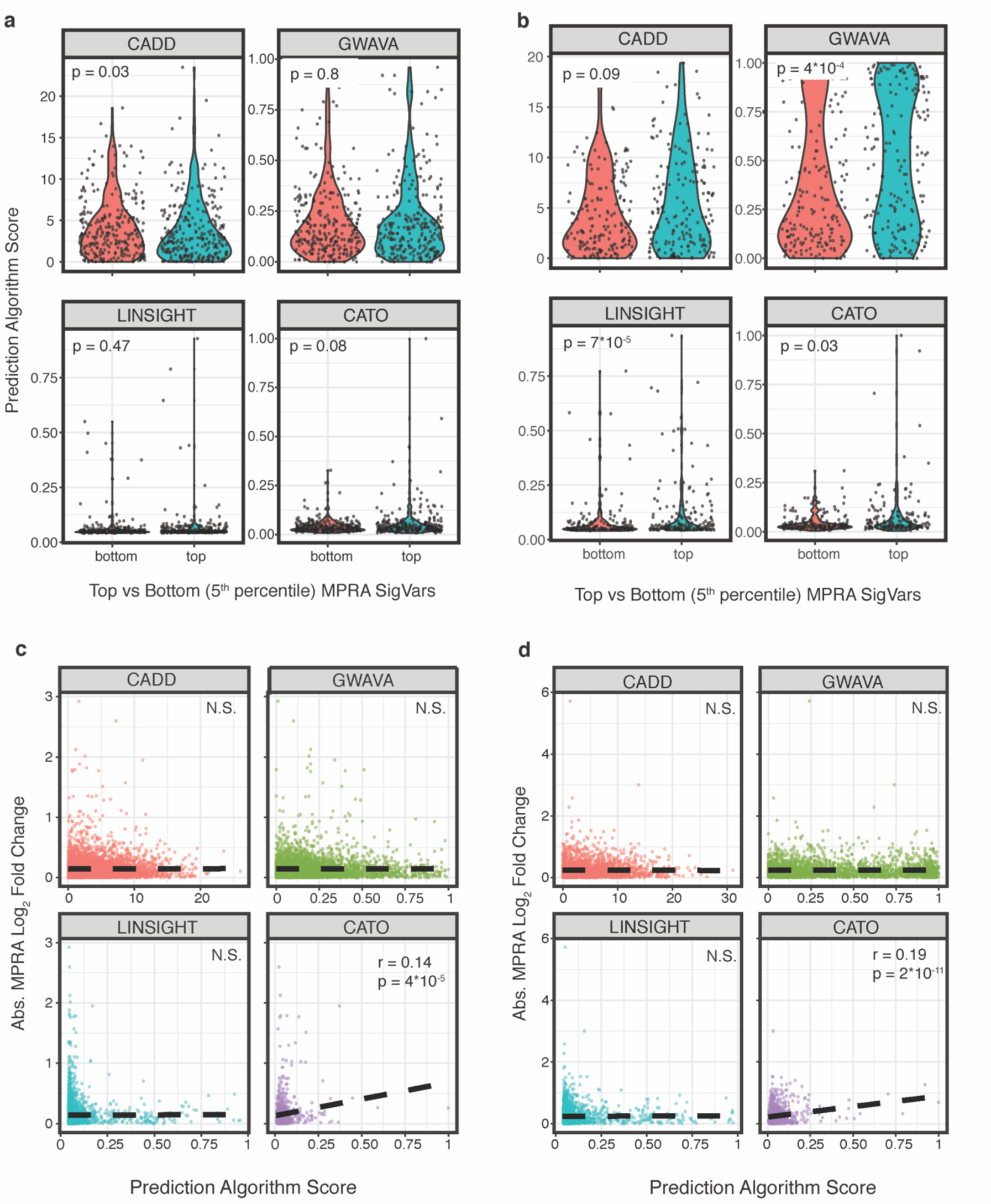
(**A-B**) All MPRA-tested variants were scored using four variant-effect prediction algorithms; LINSIGHT, CADD, GWAVA, and CATO. Violin plots show algorithm prediction scores for the top vs. bottom 5^th^ percentile of ranked variants (rank determined by MPRA allelic skew p-values) in the current study (**A**) and replication dataset (**B**). Mann-Whitney-U test. (**C-D**) When comparing all tested variants, prediction algorithm scores correlated poorly with MPRA-determined allelic skew effect sizes in both our dataset (**C**) and the replication dataset (**D**). Black dashed line = OLS regression line of best fit. Pearson’s r, all p > 0.1 (N.S.), except for CATO algorithm.

**Figure S5:**
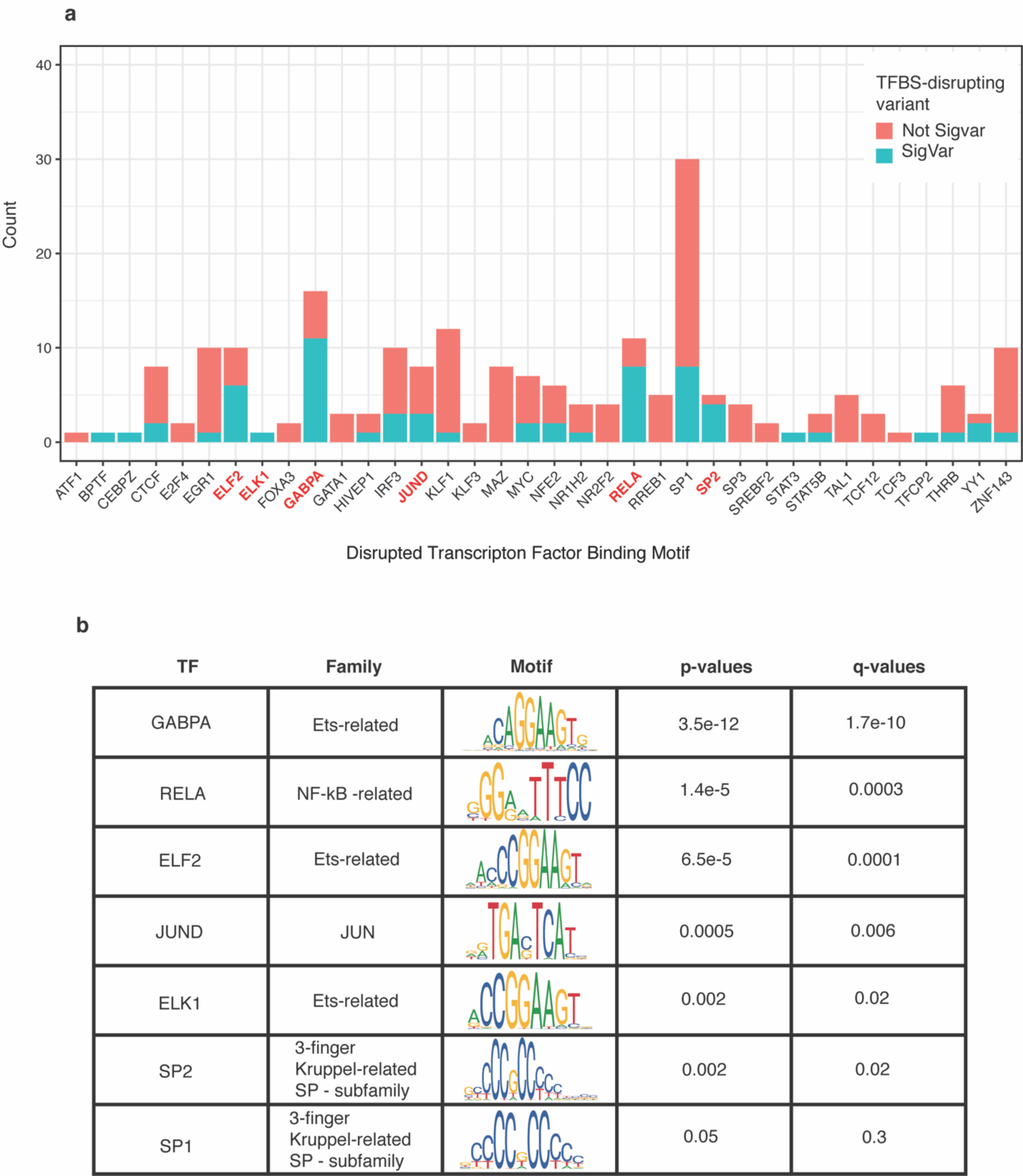
Transcription Factor Binding Site disruption is predictive of MPRA allelic skew for specific classes of TFs, including the ETS-related TF family. (**A**) Variants with significant allelic skew (SigVars) were defined at an FDR-adjusted threshold of q < 0.01 (Benjamini-Hochberg method) for variants tested in a previously published large MPRA dataset. These variants were also assessed for predicted TFBS disruption (motifbreakR; Methods) filtering for motifs corresponding to the top 200 most highly expressed TFs in K562 cells. The histogram counts the number of TFBS-disrupting variants annotated for each TF, colored by whether variants are also MPRA SigVars (blue) or not (red). TFs for which disrupted binding significantly predicts MPRA allelic skew are highlighted in red, and also described further in (**B**). Determination of a significantly increased proportion of SigVars amongst variants predicted to disrupt specific TFBSs was assessed using a one-sided binomial test, with p-values and FDR-adjusted q-values shown (Methods).

**Figure S6:**
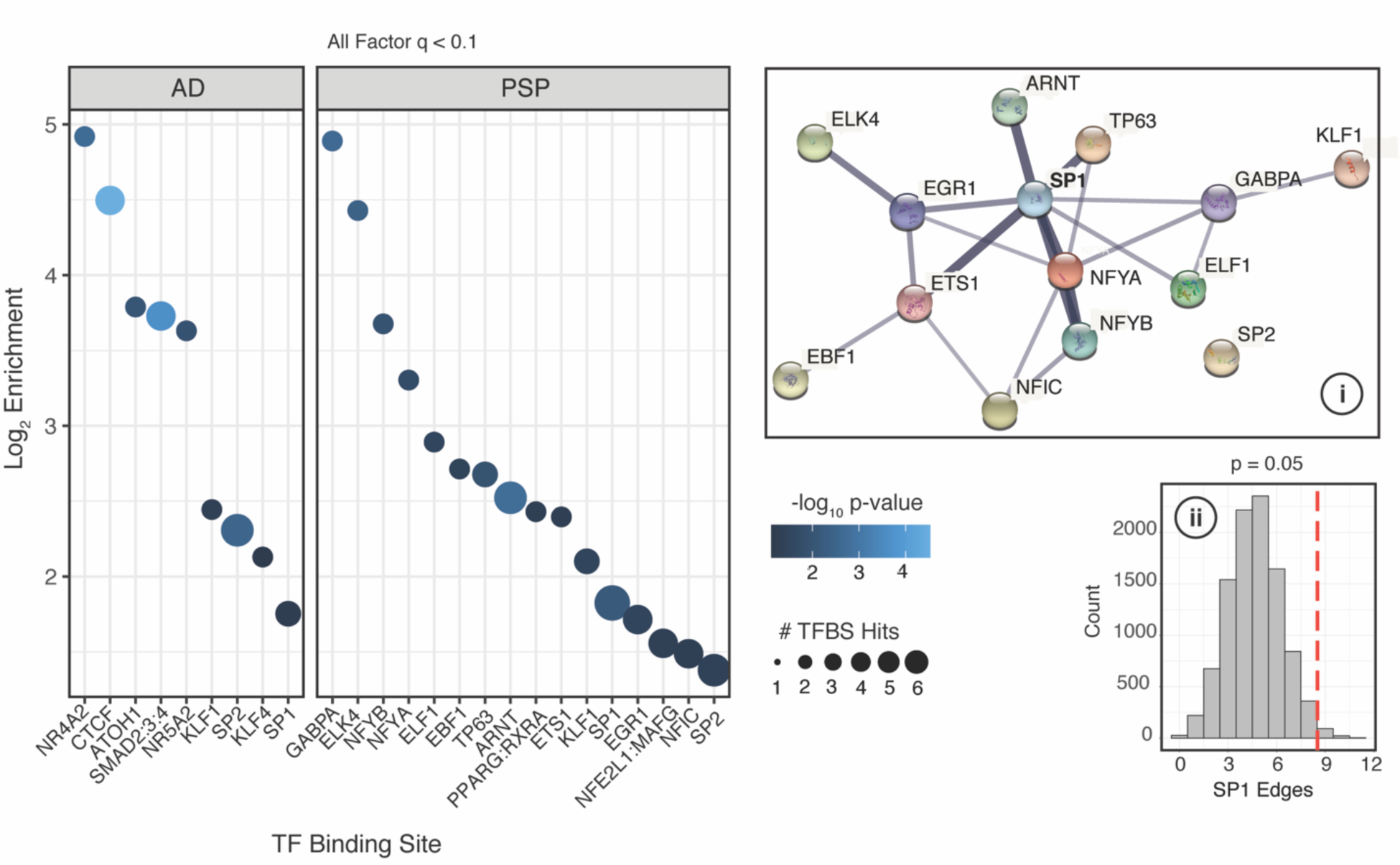
Shows the log_2_ enrichments for significantly disrupted TFBSs (FDR q < 0.1, BH method), with SigVars from AD and PSP analyzed separately (color = -log_10_ enrichment p-values, size = # disrupted TFBSs; SNPS2TFBS). Inset: i) protein-protein interaction network from the STRING database for significantly disrupted PSP-TFs. Line thickness = strength of evidence. ii) Empirical distributions for expected PPI networkSP1 - connectivity generated by resampling (Methods). Red lines = observed PSP-TF network edges.

**Figure S7:**
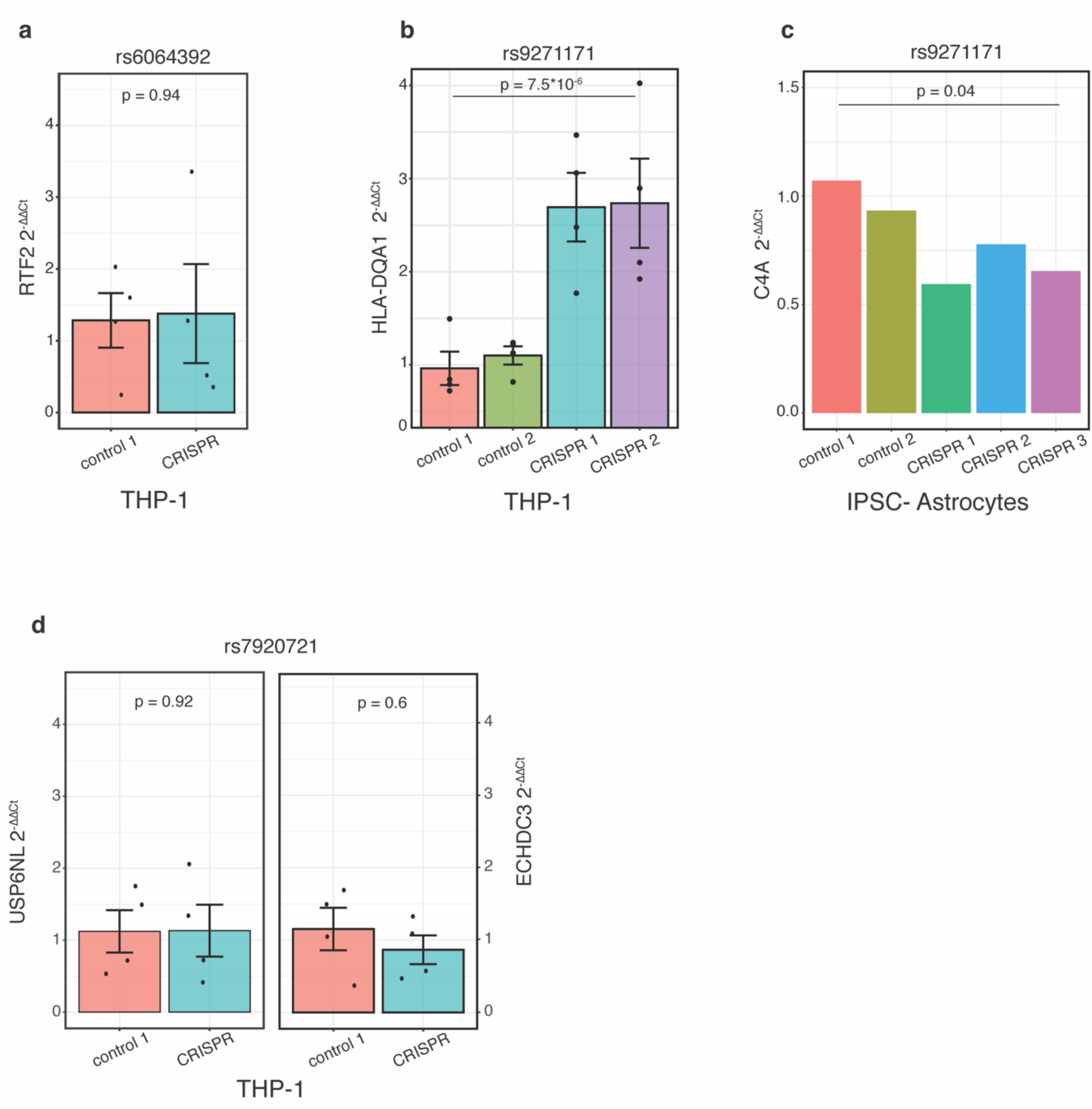
(**A**) CRISPR-mediated deletion of a small genomic region containing rs6064392 in differentiated THP-1 cells does not alter expression of *RTF2* (n=4/group; t(6) = 0.1; p = 0.94; two-tailed Student’s t-test). (**B-C**) CRISPR-mediated deletion of genomic regions containing rs9271171 significantly increases expression of *HLA-DQA1* (n=4 group, n=8/condition, t(14) = - 6.9; p = 7.5*×*10^-6^; two-tailed Student’s t-test) in differentiated THP-1 cells (**B**) and decreases expression of *C4A* in IPSC-derived astrocytes (CRISPR n = 3, control n = 2; t(3) = 3.5; p = 0.04; two-tailed Student’s t-test). (**D**) CRISPR-mediated deletion of a small genomic region containing rs7920721 in differentiated THP-1 cells does not alter expression of *USP6NL* or *ECHDC3* (n=4/group; t(6) = 0.1 and 0.6; p = 0.92 and 0.6 respectively; two-tailed Student’s t-test). Error bars = S.E.M.

**Figure S8:**
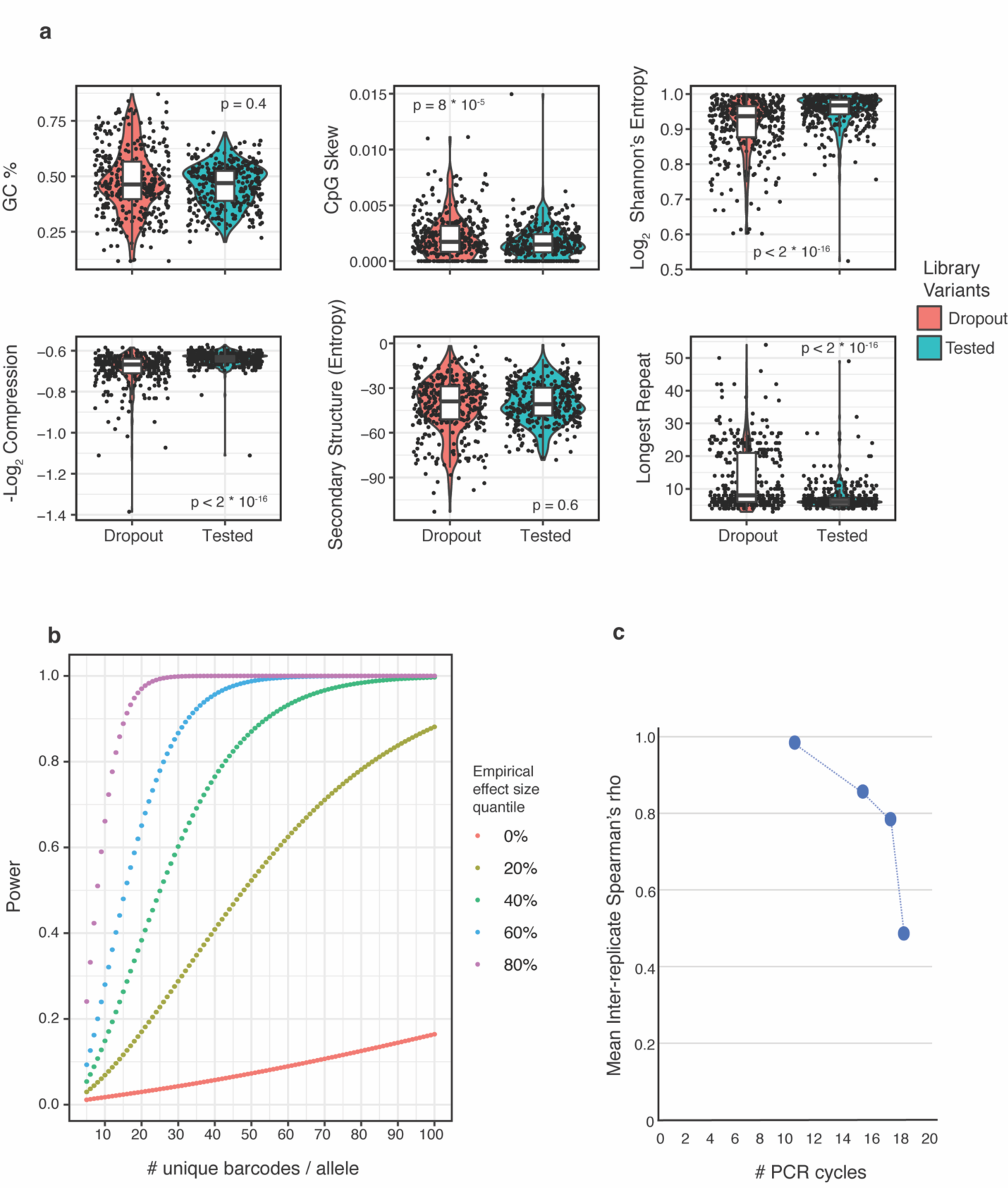
Characterization of technical features impacting MPRA performance (Supplemental Text). (**A**) Assessment of sequence-level features of the 366 variants (“Dropout”) that failed to pass quality control thresholds through both MPRA experimental stages (Methods). Violin plots display GC-content, CpG content, sequence complexity measures (-log_2_ compressibility and log_2_ Shannon’s entropy), predicted secondary structure (calculated using RNAfold), and longest mono- or di-nucleotide repeat per tested oligo for the 366 dropout variants vs. a random sample of 366 variants that passed QC (“Tested”). Dropout variants were enriched for GC content extremes (>75% or <25%), increased CpG skew, had decreased mean sequence complexity, and on average contained longer runs of nucleotide repeats (mean 13.2 vs 7.2 nucleotides) (2-tailed Mann-Whitney-U test). Predicted secondary structure did not differ for dropout variants (p > 0.05). (**B**) Simulated power analysis, using MPRA-determined empirical variance and different quantiles of empirical effect sizes, revealed that adequate power (0.8) to detect significant allelic skew at a broad range of effect-sizes is achieved at a barcode complexity of ∼40 barcodes/allele. (**C**) Plots mean correlations (Spearman’s rho) of inter-replicate allele expression for four separate MPRA experiments. Inter-replicate reproducibility is inversely proportional to the number of PCR cycles needed to amplify library mRNA before sequencing.

**Table S1.**
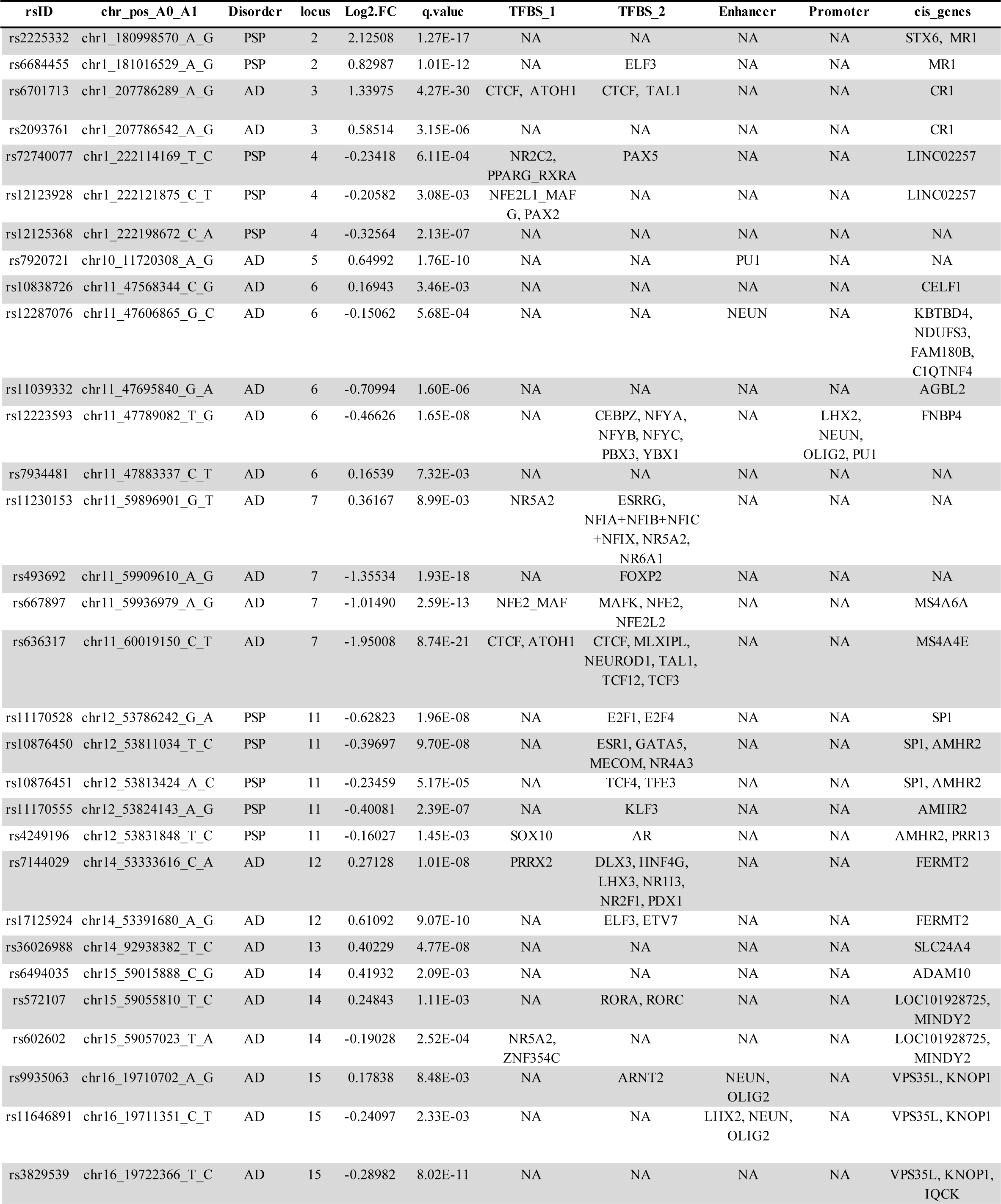
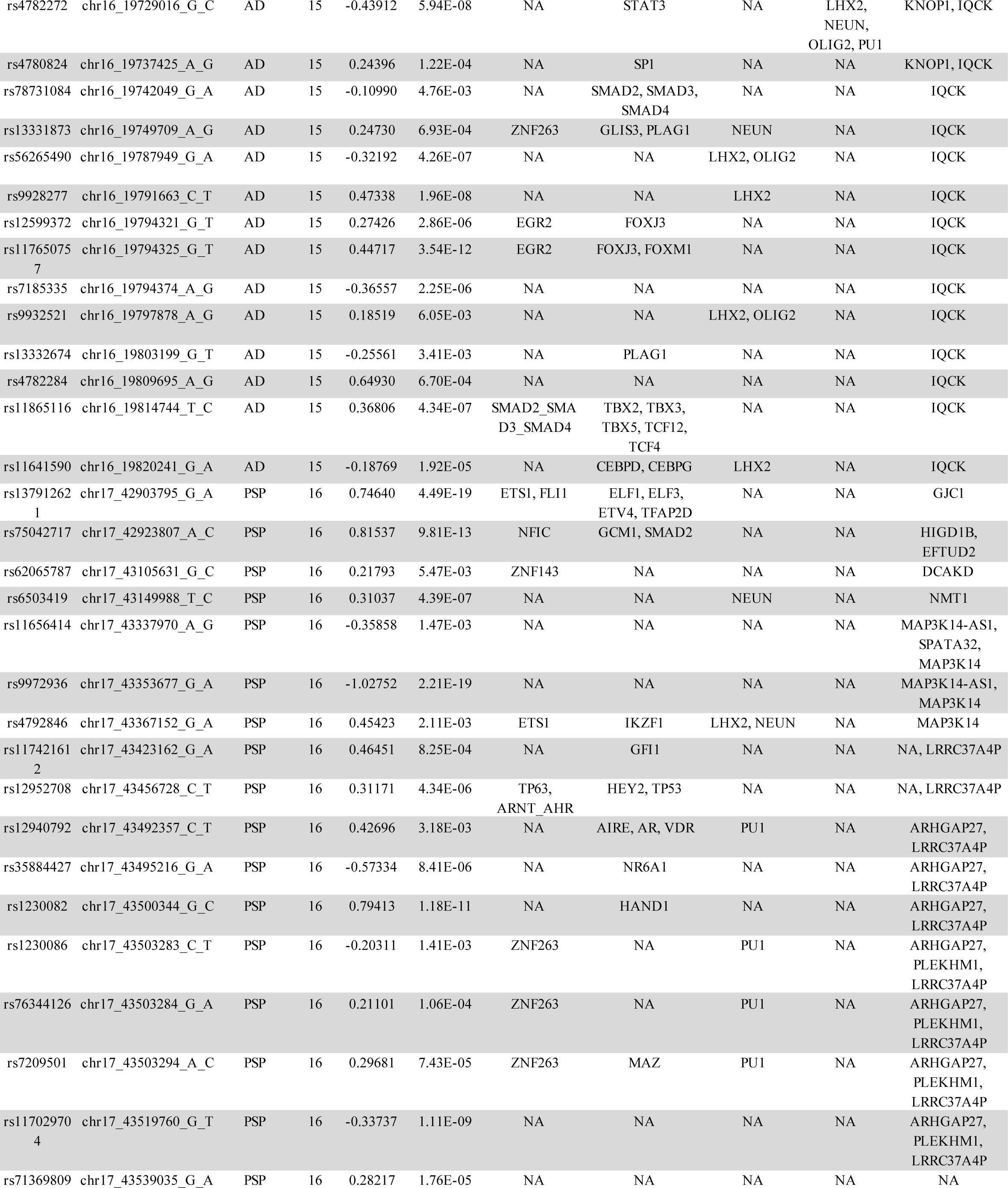
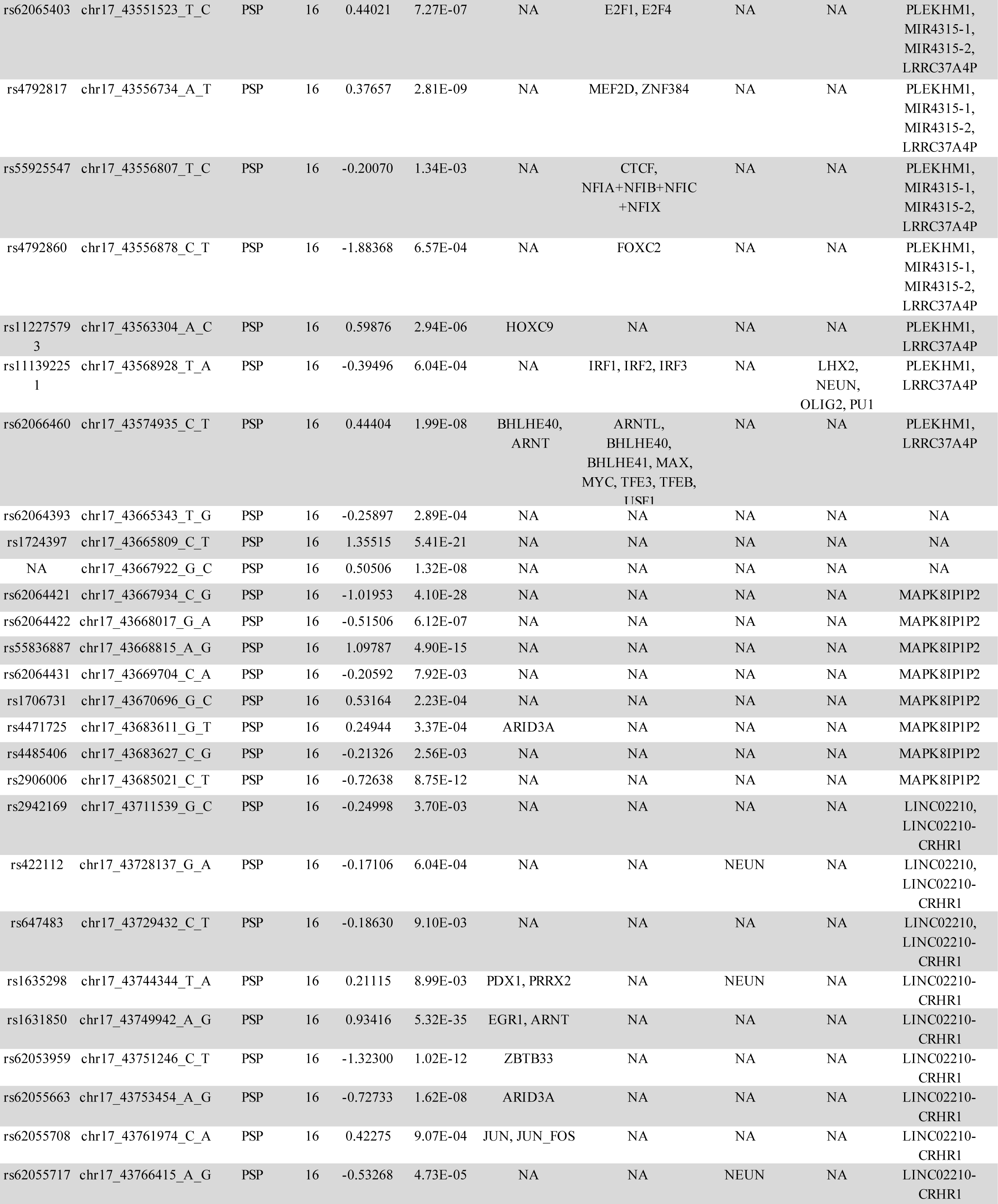
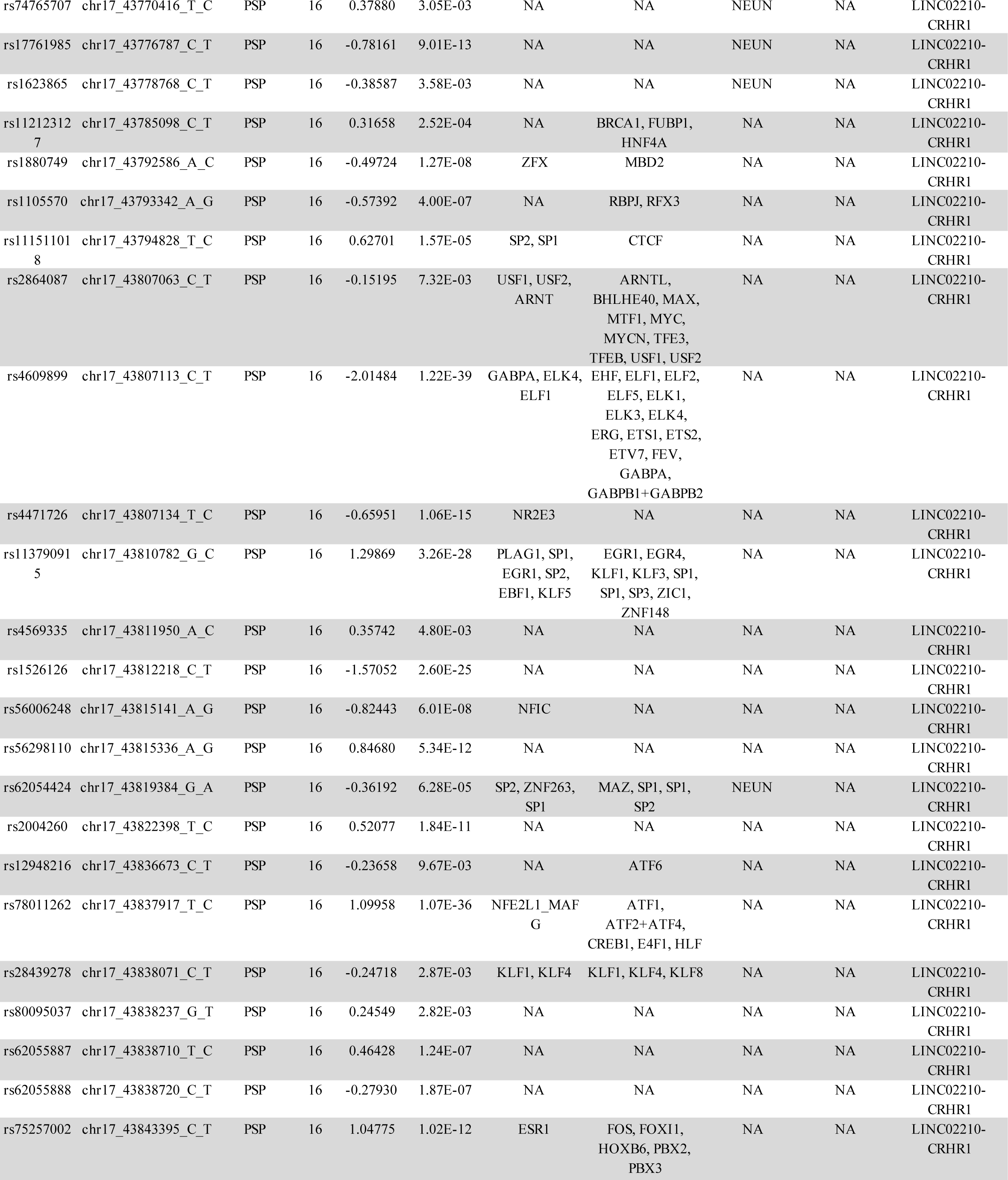
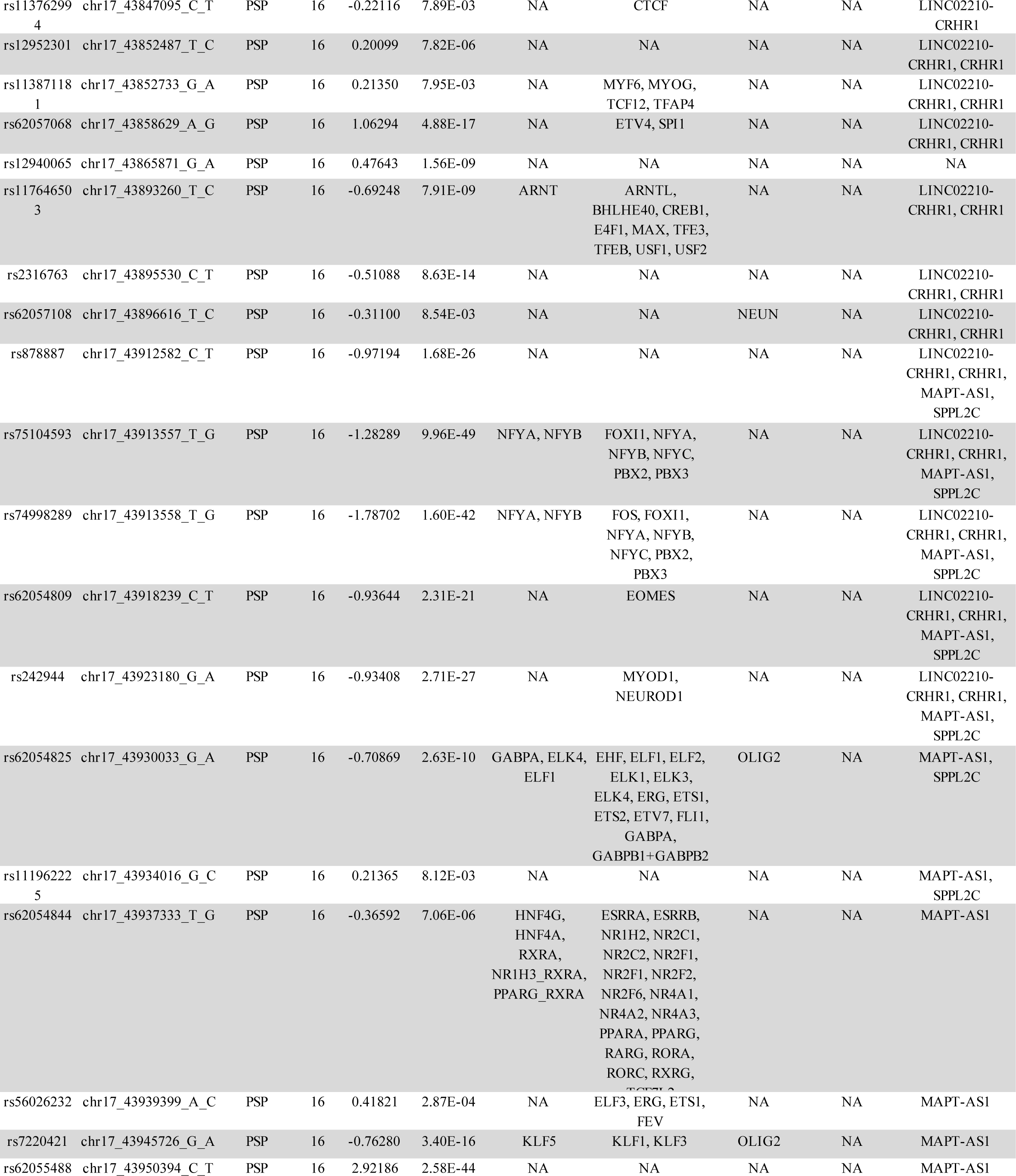
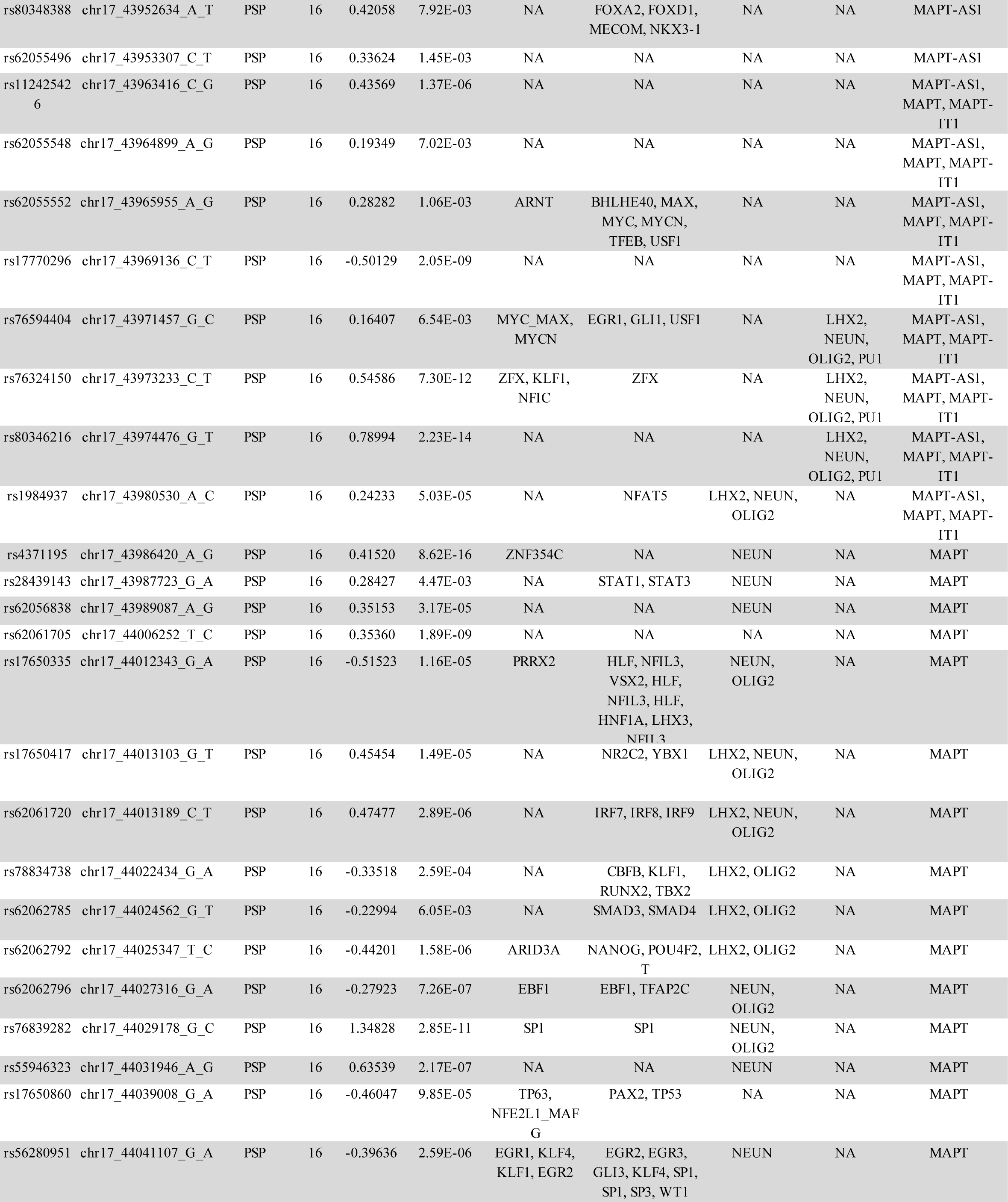
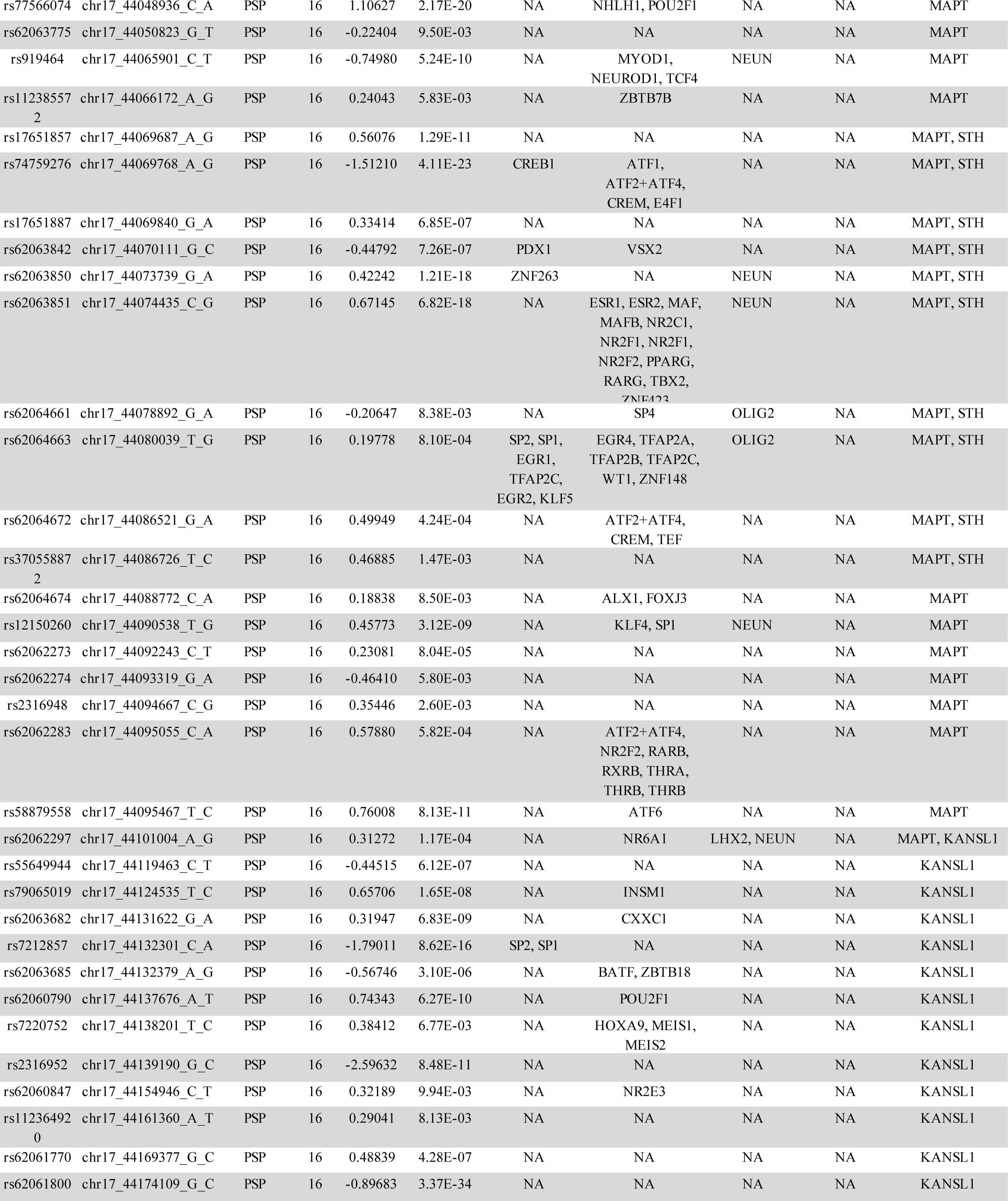
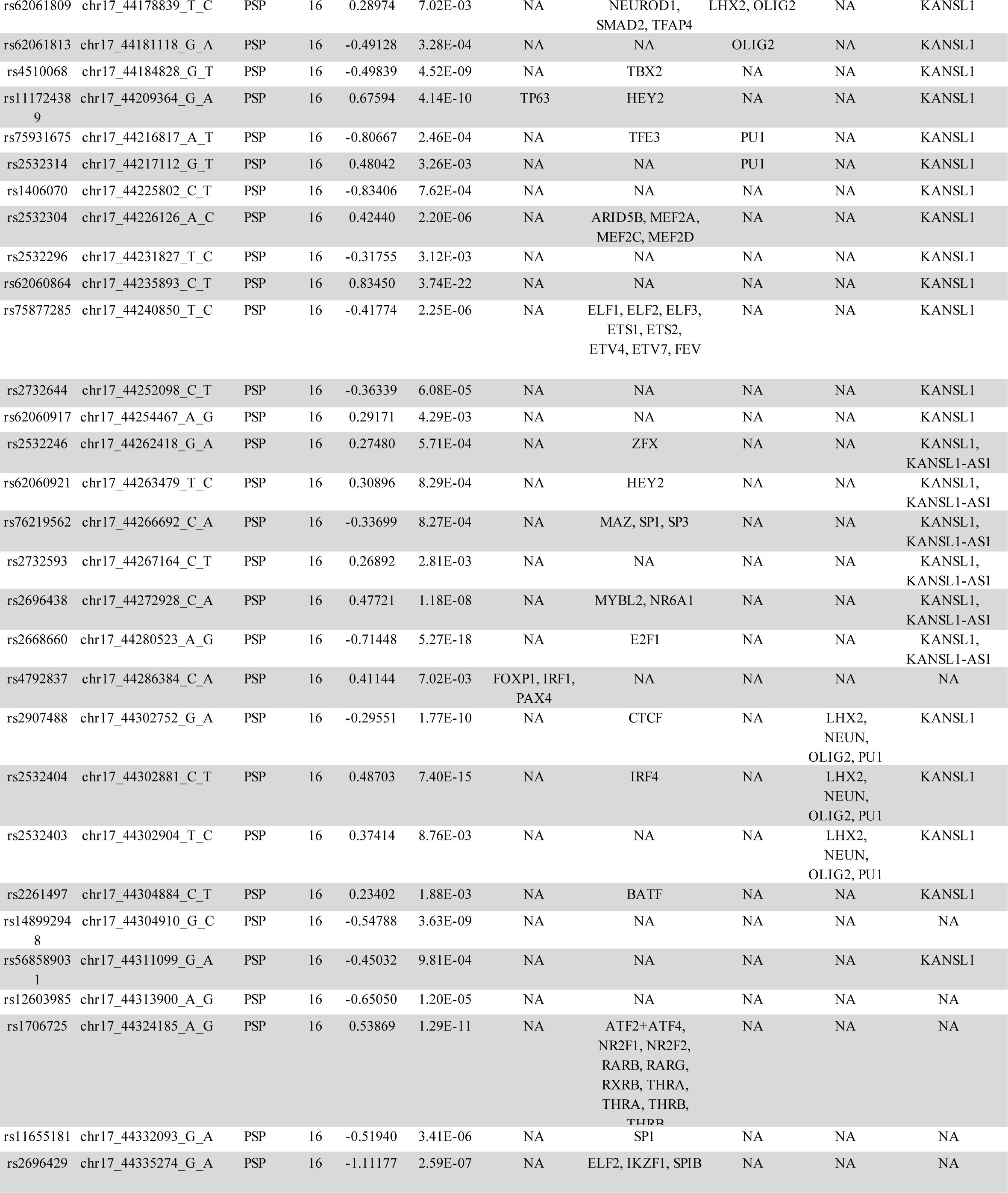
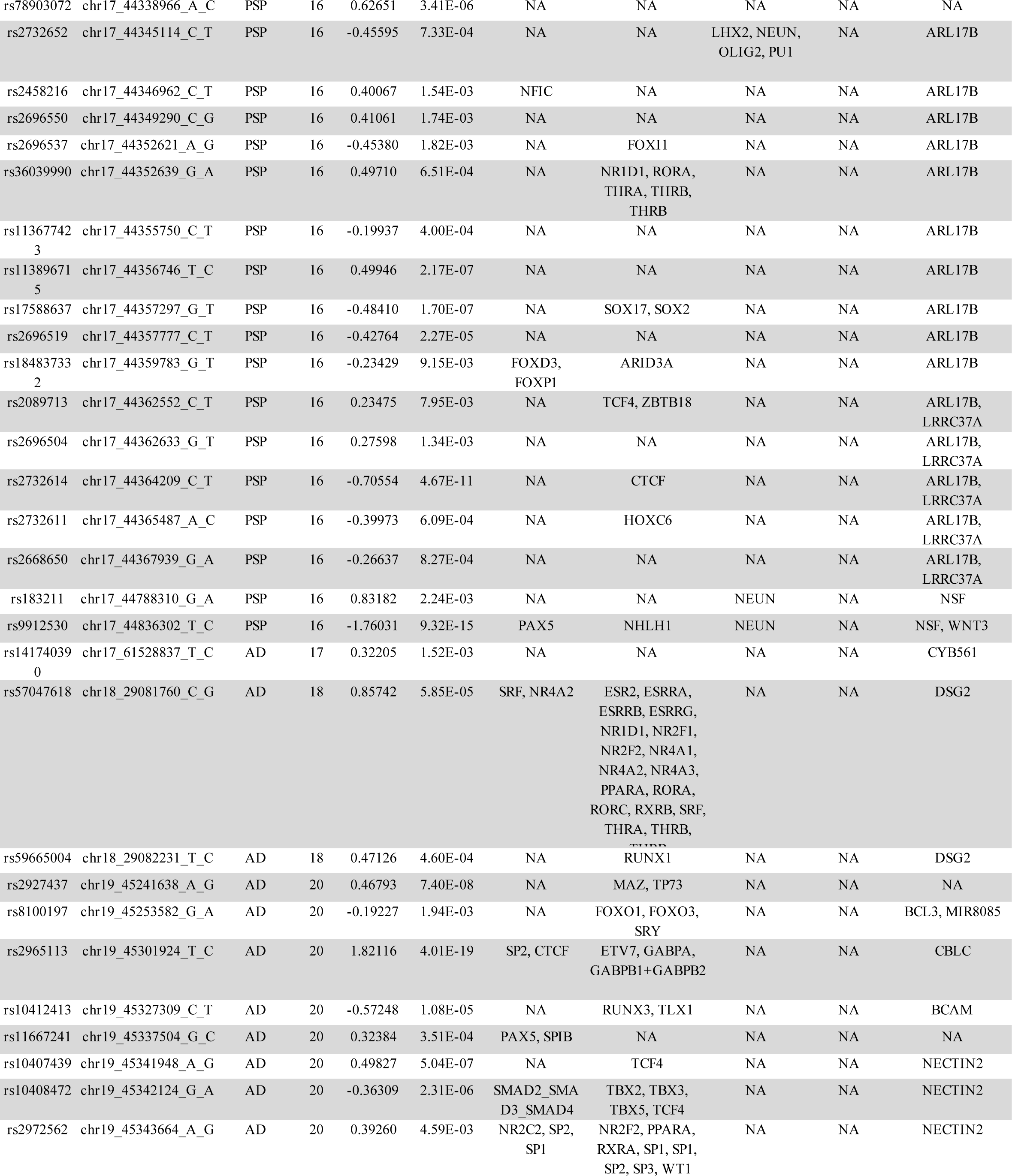
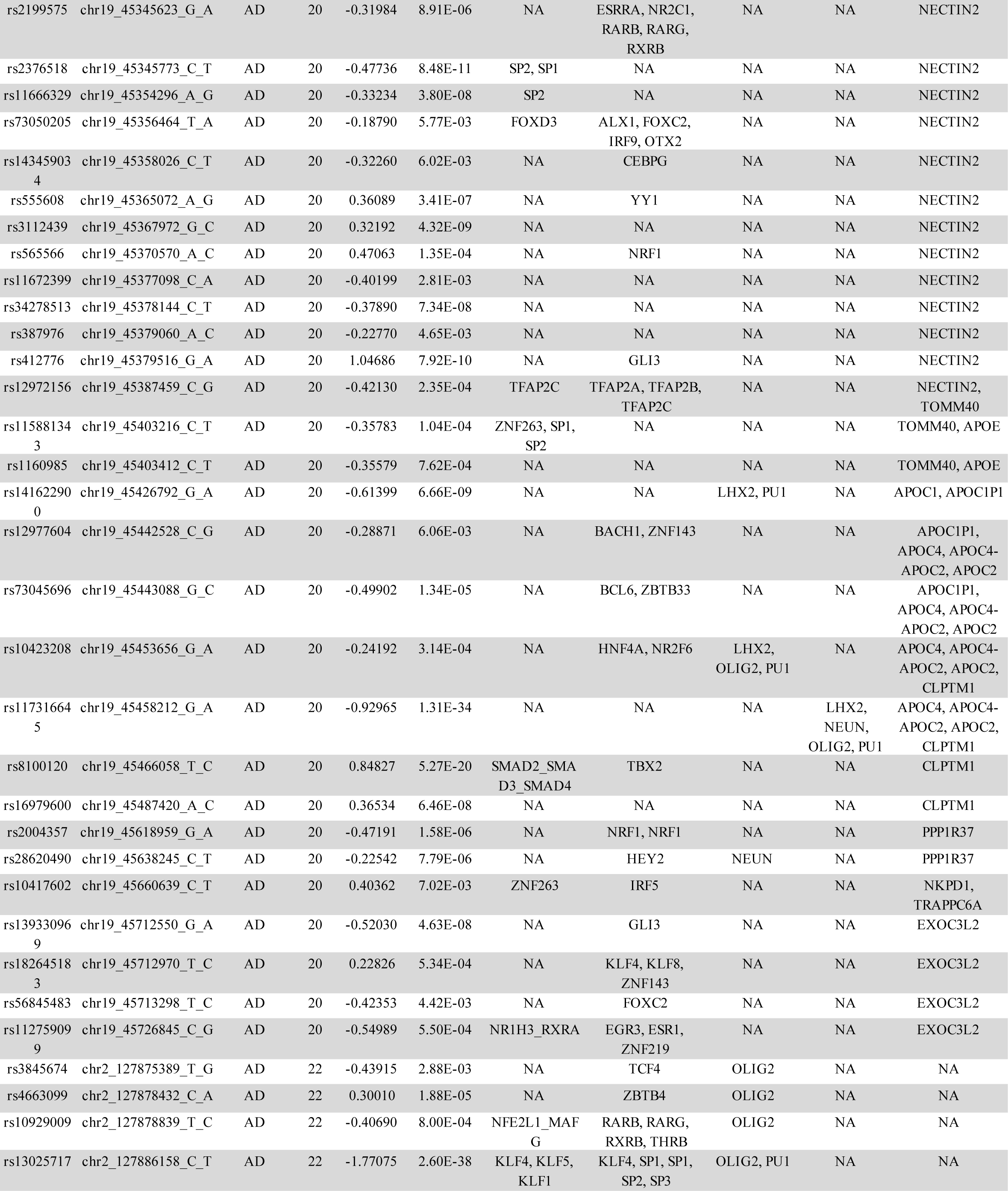
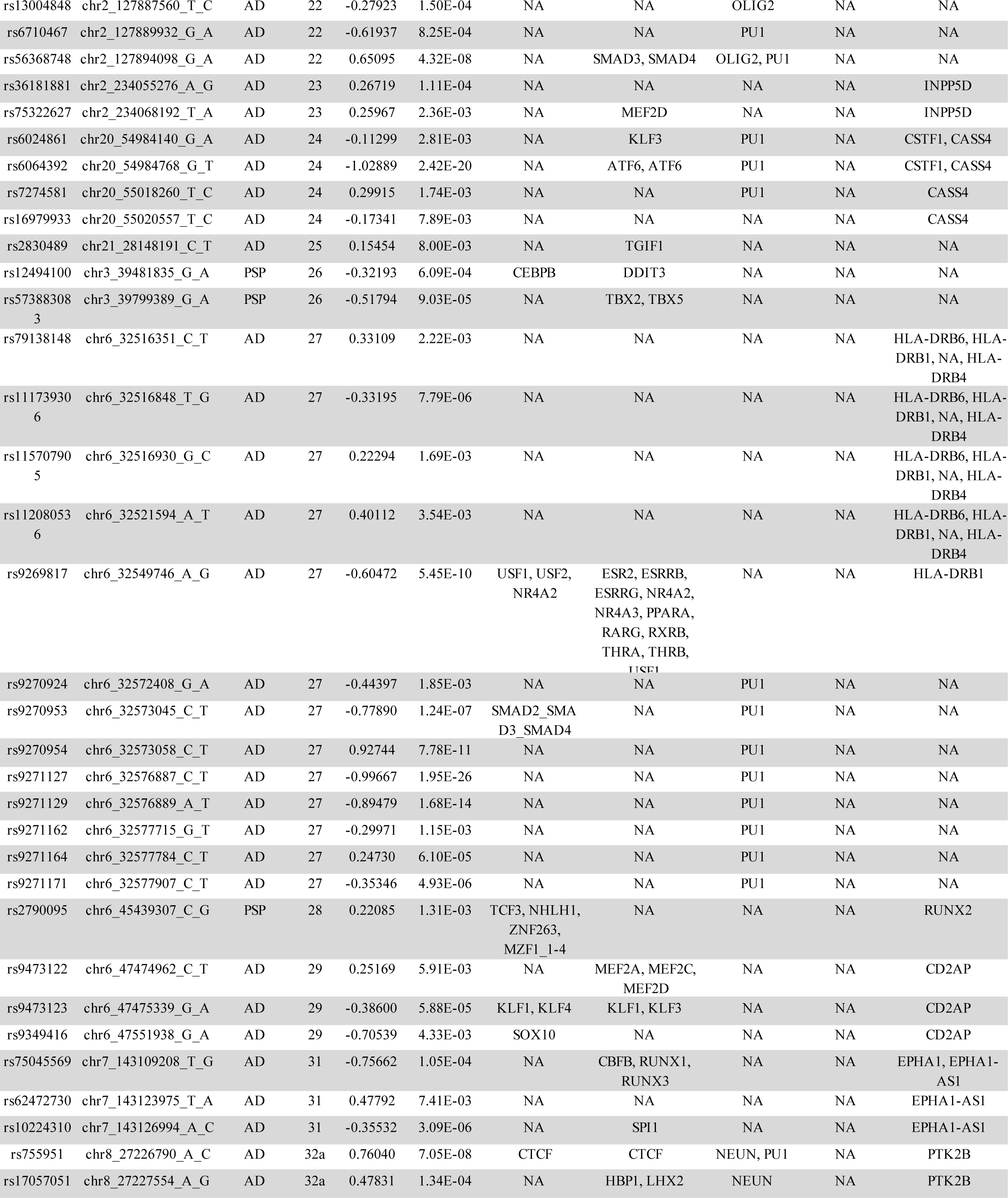
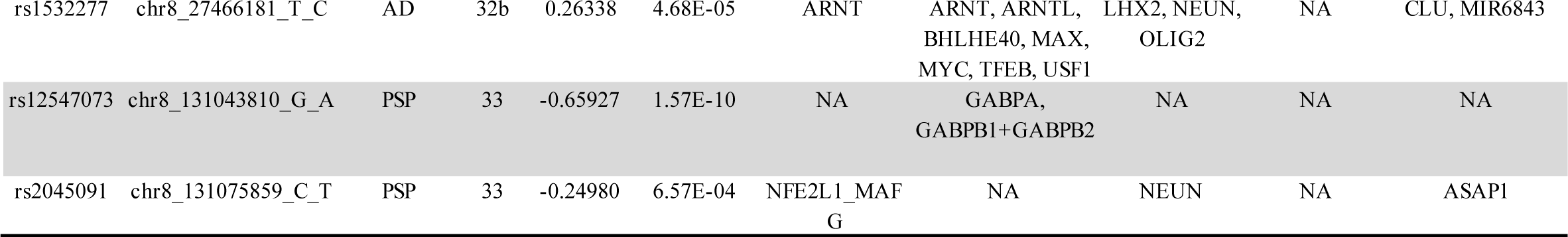
SigVars with Annotations. All 320 variants with significant allelic skew (SigVars) identified across both MPRA stages. Locus annotations are provided in Supplemental Data. SigVars are annotated with predicted TFBS-disruption using two algorithms (TFBS_1: SNPS2TFBS, TFBS_2: motifbreakR), and cell-type specific Enhancer and Promoter marks from human brain provided by Nott *et al*., 2019. Finally, variants are annotated for cis genes (gene body +/- 10 kb).

**Table S2.**
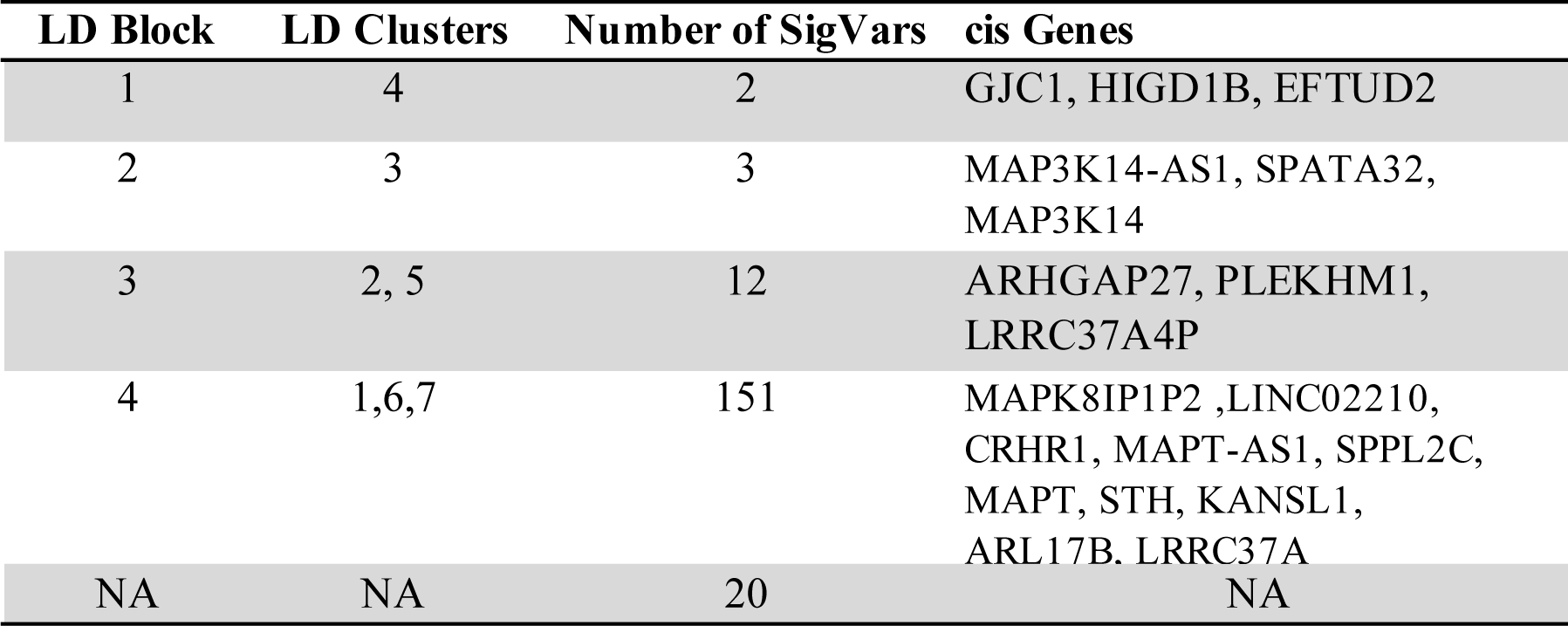
17q21.31 LD clusters, SigVars, and cis-Genes. 188 SigVars identified in the 17q21.31 region clustered by LD (CLQD function; Methods; Source Data for individual variant annotations) further organized into contiguous LD blocks. The number of SigVars per LD block and annotated cis genes (+/- 10kb) are shown.

| LD Block | LD Clusters | Number of SigVars | cis Genes |
| --- | --- | --- | --- |
| 1 | 4 | 2 | GJC1, HIGD1B, EFTUD2 |
| 2 | 3 | 3 | MAP3K14-AS1, SPATA32, MAP3K14 |
| 3 | 2, 5 | 12 | ARHGAP27, PLEKHM1, LRRC37A4P |
| 4 | 1,6,7 | 151 | MAPK8IP1P2 ,LINC02210, CRHR1, MAPT-AS1, SPPL2C, MAPT, STH, KANSL1, ARL17B, LRRC37A |
| NA | NA | 20 | NA |

**Supplemental Table 3:** SigVars per gene. Genes for which there was an MPRA SigVar within +/- 10 kb of the gene body. “SigVar Number” is the number SigVars/gene. Some SigVars are annotated to multiple nearby genes. Genes for which there was an MPRA SigVar within +/- 10 kb of the gene body. “SigVar Number” is the number SigVars/gene. Some SigVars are annotated to multiple nearby genes.

| chr | Gene | Sigvar Number |
| --- | --- | --- |
| chr15 | ADAM10 | 1 |
| chr11 | AGBL2 | 1 |
| chr12 | AMHR2 | 4 |
| chr19 | APOC1 | 1 |
| chr19 | APOC1P1 | 3 |
| chr19 | APOC2 | 4 |
| chr19 | APOC4 | 4 |
| chr19 | APOC4-APOC2 | 4 |
| chr19 | APOE | 2 |
| chr17 | ARHGAP27 | 7 |
| chr17 | ARL17B | 15 |
| chr8 | ASAP1 | 1 |
| chr19 | BCAM | 1 |
| chr19 | BCL3 | 1 |
| chr11 | C1QTNF4 | 1 |
| chr20 | CASS4 | 4 |
| chr19 | CBLC | 1 |
| chr6 | CD2AP | 3 |
| chr11 | CELF1 | 1 |
| chr19 | CLPTM1 | 4 |
| chr8 | CLU | 1 |
| chr1 | CR1 | 2 |
| chr17 | CRHR1 | 11 |
| chr20 | CSTF1 | 2 |
| chr17 | CYB561 | 1 |
| chr17 | DCAKD | 1 |
| chr18 | DSG2 | 2 |
| chr17 | EFTUD2 | 1 |
| chr7 | EPHA1 | 1 |
| chr7 | EPHA1-AS1 | 3 |
| chr19 | EXOC3L2 | 4 |
| chr11 | FAM180B | 1 |
| chr14 | FERMT2 | 2 |
| chr11 | FNBP4 | 1 |
| chr17 | GJC1 | 1 |
| chr17 | HIGD1B | 1 |
| chr6 | HLA-DRB1 | 5 |
| chr6 | HLA-DRB4 | 4 |
| chr6 | HLA-DRB6 | 4 |
| chr2 | INPP5D | 2 |
| chr16 | IQCK | 15 |
| chr17 | KANSL1 | 37 |
| chr17 | KANSL1-AS1 | 6 |
| chr11 | KBTBD4 | 1 |
| chr16 | KNOP1 | 5 |
| chr17 | LINC02210 | 3 |
| chr17 | LINC02210-CRHR1 | 45 |
| chr1 | LINC02257 | 2 |
| chr15 | LOC101928725 | 2 |
| chr17 | LRRC37A | 5 |
| chr17 | LRRC37A4P | 16 |
| chr17 | MAP3K14 | 3 |
| chr17 | MAP3K14-AS1 | 2 |
| chr17 | MAPK8IP1P2 | 8 |
| chr17 | MAPT | 45 |
| chr17 | MAPT-AS1 | 21 |
| chr17 | MAPT-IT1 | 8 |
| chr15 | MINDY2 | 2 |
| chr17 | MIR4315-1 | 4 |
| chr17 | MIR4315-2 | 4 |
| chr8 | MIR6843 | 1 |
| chr19 | MIR8085 | 1 |
| chr1 | MR1 | 2 |
| chr11 | MS4A4E | 1 |
| chr11 | MS4A6A | 1 |
| chr11 | NDUFS3 | 1 |
| chr19 | NECTIN2 | 16 |
| chr19 | NKPD1 | 1 |
| chr17 | NMT1 | 1 |
| chr17 | NSF | 2 |
| chr17 | PLEKHM1 | 11 |
| chr19 | PPP1R37 | 2 |
| chr12 | PRR13 | 1 |
| chr8 | PTK2B | 2 |
| chr6 | RUNX2 | 1 |
| chr14 | SLC24A4 | 1 |
| chr12 | SP1 | 3 |
| chr17 | SPATA32 | 1 |
| chr17 | SPPL2C | 7 |
| chr17 | STH | 10 |
| chr1 | STX6 | 1 |
| chr19 | TOMM40 | 3 |
| chr19 | TRAPPC6A | 1 |
| chr16 | VPS35L | 3 |
| chr17 | WNT3 | 1 |

Data S1: Supplemental Data

This file provides summary statistics for all variants tested in MPRA stages 1 and 2

Data S2: Source Data

This file provides raw data and computed scores used for downstream analyses and figure construction. Data are organized by relevant figure.

Data S3: Supplemental Information - Methods

This file provides additional Methods data including sequences, primers, and accessions.

